# Ischemia-Induced Post-Translational Modifications of GLT-1 Mediate Aberrant Trafficking and Impaired Glutamate Uptake

**DOI:** 10.64898/2026.03.13.711603

**Authors:** Simran Kaur Gill, Katelyn Louise Reeb, Max Kroll, Ole V. Mortensen, Andréia Cristina Karklin Fontana

**Author notes:** **Corresponding author:** Andréia C. K. Fontana, Department of Pharmacology and Physiology, Drexel University College of Medicine, Philadelphia, PA, 19102, United States.

## Abstract

Glutamate transporters are essential for maintaining CNS homeostasis by clearing extracellular glutamate after synaptic transmission. Dysregulation of these transporters contributes to excitotoxicity in numerous neurological disorders, including ischemic stroke, making them a promising therapeutic target. However, the regulatory response of these transporters following ischemic insult remains poorly defined. In this work, using a model of oxygen-glucose deprivation in primary glia cultures, we report aberrant trafficking of the astrocytic glutamate transporter GLT-1 following ischemic insult. This response is characterized by increased transporter internalization and degradation, accompanied by reduced glutamate uptake capacity. Focusing on post-translational modifications (PTMs), we found that GLT-1 ubiquitination is markedly increased after ischemic insult and contributes to transporter internalization. Importantly, disrupting this ubiquitination interaction through mutation of C-terminal GLT-1 lysine residues restores GLT-1 surface expression and rescues glutamate uptake capacity through preventing early endosome 1 (EEA1)-mediated internalization. Additionally, we report that inhibition of C-terminal GLT-1 PTMs confers neuroprotection following ischemic insult in organotypic rat brain slices. Together, these findings demonstrate that ischemia-induced dysregulation of GLT-1 trafficking plays a critical role in impaired glutamate clearance and cellular recovery, highlighting GLT-1 ubiquitination as a potential therapeutic target for ischemic injury.

## 1 Introduction

Glutamate, the most abundant neurotransmitter in the central nervous system (CNS), plays a central role in regulating neuronal communication, synaptic plasticity, and overall neural homeostasis (Danbolt, 2001). Excitatory glutamatergic signaling is tightly balanced by its inhibitory counterpart, γ-aminobutyric acid (GABA), and together these systems preserve proper brain function (Zhou & Danbolt, 2014). Even small disruptions in the levels of either neurotransmitter can disturb this homeostatic equilibrium, leading to profound consequences for CNS signaling (Lipton & Rosenberg, 1994). In particular, dysregulated glutamate signaling has been implicated in a range of neurological and neuropsychiatric disorders (O’Donovan et al., 2017; McGrath et al., 2022) including stroke (Choi, 1988; Bramlett & Dietrich, 2004; Patel & McMullen, 2017), neuropathic pain (Osikowicz et al., 2013; Temmermand et al., 2022), epilepsy (Choi, 1988; Tanaka et al., 1997; Green et al., 2021) and traumatic brain injury (Rao et al., 2001; Ikematsu et al., 2002).

Stroke remains a leading cause of death and long-term disability worldwide (Doyle et al., 2008; de Havenon & Sheth, 2025). Ischemic stroke, the most prevalent subtype of stroke, results from inadequate cerebral blood flow that restricts the delivery of oxygen and glucose, and most commonly is due to vascular occlusion. Deprivation of these essential substrates leads to neuronal cell death (Puig et al., 2018), primarily through glutamate-induced excitotoxicity, a neurotoxic process initiated by excess glutamate release and overactivation of ionotropic glutamate receptors (Choi, 1992). This hyper-glutamatergic state leads to excessive calcium influx, ultimately activating neuronal cell death pathways (Goldberg & Choi, 1993; Cross et al., 2010).

An important mediator of glutamatergic homeostasis is the family of excitatory amino acid transporters (EAATs), which clear glutamate from the synaptic cleft. They are comprised of five excitatory amino acid transporter (EAAT) subtypes, EAAT1–5 (in rodents, EAAT1, EAAT2, and EAAT3 are commonly referred to as GLAST, GLT-1, and EAAC1, respectively). EAAT1 and EAAT2 are primarily located on astrocytes, with EAAT2 responsible for over 90% of glutamate clearance in the CNS (Suchak et al., 2003). Therefore, EAAT2 (hereafter referred to as GLT-1) is crucial for synaptic glutamate clearance and limiting excitotoxic damage.

Although EAATs are known to be critical mediators of glutamate homeostasis after ischemic insult (Rao et al., 2001; Chu et al., 2007; Weller et al., 2008), the regulation of these transporters is not well defined. Specifically, the signaling pathways that govern changes in glutamate transporter activity and expression over the course of ischemia have yet to be determined. Previous studies have shown that GLT-1 levels decrease following ischemia in rats (J. C. Chen et al., 2005), and that GLT-1 knockdown worsens neuronal damage while increasing mortality rates (Rao et al., 2001). Additionally, strategies to enhance GLT-1 expression reduce infarct volume and improve neurological deficits (Krzyzanowska et al., 2017). Severe ischemic insults can also lead to a reversal in glutamate transport, further contributing to extracellular glutamate accumulation (Phillis & O’Regan, 1996; Grewer et al., 2008; Wang et al., 2013). Despite these observations, the mechanisms regulating GLT-1 trafficking and surface availability in response to ischemic insult are not well defined.

Expression levels of GLT-1 are tightly regulated at multiple levels, including transcriptional, translational, and post-translational mechanisms. At the transcriptional level, several regulatory proteins influence GLT-1 expression. For example, Ying Yang 1 (YY1) has been identified as a negative regulator, while other transcription factors such as NF-κB and N-Myc can either enhance or repress GLT-1 expression depending on cellular context and external stimuli (Martinez-Lozada et al., 2016). At the post-transcriptional and translational level, GLT-1 is regulated through diverse mechanisms, including protein–protein interactions via PDZ domain-containing proteins (Bassan et al., 2008), and post-transcriptional regulation by microRNAs in response to cellular stress or injury (Zumkehr et al., 2018; Meng et al., 2021). Among these mechanisms, post-translational modifications (PTMs) play a key role in regulating GLT-1 trafficking and functionality (Peterson & Binder, 2019). In physiological states, several PTMs such as the ubiquitination and SUMOylation drive basal transporter turnover and cellular compartmentalization, which both primarily occur at lysine residues of the C-terminal domain (Martínez-Villarreal et al., 2012; Foran et al., 2014). However, dysregulation of GLT-1 PTMs has been implicated in abnormal transporter localization and expression across multiple neurological disorders. For example, increased GLT-1 ubiquitination and subsequent degradation has been reported in a mouse model of Parkinson’s disease (Zhang et al., 2017), whereas reduced transporter palmitoylation in Huntington’s disease lead to a decrease in glutamate uptake (Huang et al., 2010). However, the role of PTMs in governing GLT-1 localization and trafficking in ischemic conditions is not completely understood.

In this study, we investigated the role of PTMs in regulating GLT-1 trafficking following ischemic insult. We demonstrate that ischemia induces ubiquitination-dependent internalization and degradation of GLT-1 in response to ischemic insult in glia cultures and brain slices. Furthermore, preventing C-terminal PTMs restores GLT-1 surface expression and functional glutamate uptake, thereby conferring neuroprotection.

## 2 Methods

### 2.1 Animals

Sprague-Dawley dams with their pups (ordered at postnatal age day 2) were obtained from Charles River Laboratory (Malvern, PA, USA). Animals were housed in the Animal Facility at Drexel University College of Medicine that is accredited to the Association for Assessment and Accreditation of Laboratory Animal Care (AAALAC). A total of 45 dams and 135 pups (male and female) were used for this study. Animals were provided with access to food and water *ad libitum*. Animal housing and experimental procedures were approved by the Drexel University College of Medicine Institutional Animal Care and Use Committees (IACUC) under protocols #209-7 and #LA-23-749.

### 2.2 Materials

For COS-7 cells, primary glia cultures, and organotypic slice cultures (OSCs), Dulbecco’s modified Eagle’s medium (DMEM) with and without glucose (cat: 11995065 and cat: 11966025), 2.5% trypsin (cat: 15090-046), gentamicin (cat: 15750060), 96-well poly-L-lysine precoated plates (cat: 152039), and Countess^TM^ III FL cell counter (cat: AMQAF2000) were purchased from ThermoFisher Scientific (Waltham, MA, USA). Penicillin-streptomycin solution (cat: 30-002-CI) was purchased from Corning (Corning, NY, USA). Deoxyribonuclease I (cat: D5025-150KU) and poly-L-lysine (cat: P1274) were purchased from Sigma Aldrich (St. Louis, MO, USA). Heat inactivated fetal bovine serum (FBS) used for primary cultures (cat: SH3007003HI) was obtained from Hyclone (Logan, UT, USA), while FBS used for COS-7 cell culture (cat: 89510-186) was obtained from Avantor (Radnor, PA, USA). Millicell membrane inserts (cat: PICM03050) were purchased from EMD Millipore (Burlington, MA, USA). B27 (cat: 17504-044) and N2 (cat: 17502-048) supplements were purchased from ThermoFisher Scientific.

All lentivirus vectors used in this study were purchased from Vector Builder (Chicago, IL, USA). The transfection reagent DharmaFECT 3 was purchased from Horizon (Cambridge, UK). For pharmacological treatments, phorbol 12-myristate 13-acetate (PMA) was purchased from Sigma Aldrich (cat: 524400). TFB-TBOA (cat: 2532), UCPH-101 (cat: 3490), WAY 213613 (cat: 2652), and MK-801 (cat: 0924) were purchased from Tocris Bioscience (Bristol, UK). MG-132 (cat: HY-1325) Bafilomycin (cat: HY-100558), TAK-243 (cat: HY-100487) and Dynasore (cat: HY-15304) were purchased from MedChem Express (Monmouth Junction, NJ). For oxygen-glucose deprivation (OGD), the hypoxic chamber was obtained from Billups-Rothenberg Inc. (San Diego, CA, USA).

For glutamate uptake assays, L-[3,4-^3^H]-Glutamic Acid (specific activity 50.8 Ci/mmol, hereafter referred to as L-^3^H-glutamate) was purchased from Revvity (Waltham, MA, USA). TFB-TBOA (cat: 25-321-0) was obtained from ThermoFisher. The transfection reagent TransIT-LT1 was purchased from Mirus Bio LLC (Madison, WI, USA). EcoLite (+) scintillation cocktail was purchased from MP Biomedicals (Irvine, CA, USA). A 96-well plate washer was purchased from BioTek (Winooski, Vermont, USA), and the Microplate Scintillation and Luminescence Counter was purchased from Wallac (Shelton, CT, USA). The LS 6500 counter for 24 well plates was from Beckman Coulter, Brea, CA.

The antibodies used in this study, including their sources and concentrations, are listed in Table 1.

**Table 1:** Antibodies, sources, and dilutions used in this study.

| Antibody | Host | Source | Product cat.<br>Number and RRID | Application and<br>dilution |
| --- | --- | --- | --- | --- |
| *Primary antibodies* |  |  |  |  |
| GLT-1 | Rabbit | Gifted (NIH): targeted against C- terminal domain | N/A | Western Blot (WB)<br>1:10,000 |
|  |  |  |  | Co-IP<br>0.04 µg/µl |
| GLAST | Rabbit | Alomone Labs (Jerusalem, Israel) | AGC-021 | WB<br>1:500 |
|  |  |  |  | Immunocytochemistry (ICC)<br>1:100 |
| MAP-2 | Rabbit | EMD Millipore (Burlington, MA, USA) | AB5622ICC | ICC<br>1:1000 |
| IBA1 | Guinea Pig | Synaptic Systems (Goettingen, Germany) | 234 308 | ICC<br>1:500 |
| GFAP | Mouse | EMD Millipore (Burlington, MA, USA) | MAB360 | ICC<br>1:400 |
| Flag | Mouse | ThermoFisher (Waltham, MA, USA) | MA1-91878<br>RRID: AB_1957945 | WB<br>1:300 |
|  |  |  |  | ICC<br>1:300 |
|  |  |  |  | Co-IP<br>3µg |
| Early Endosome Antigen 1 (EEA1) | Rabbit | Cell Signaling (Danvers, MA) | 3288S<br>RRID: AB_2096811 | ICC<br>1:100 |
|  | Mouse | Becton Dickinson, BD (Franklin Lakes, NJ) | 610456<br>RRID: AB_397829 | ICC<br>1:200 |
| SUMO-1 | Mouse | Santa Cruz Biotechnology (Dallas, TX) | Sc-5308<br>RRID: AB_628300 | WB<br>1:500 |
|  |  |  |  | Co-IP<br>0.02 µg/µl |
| IgG | Rat | Sigma Aldrich<br>RRID:AB_3740246 | NI04 | Co-IP<br>0.02 µg/µl |
| P4D1 (Ubiquitin) | Mouse | Santa Cruz Biotechnology | Sc-8017<br>RRID: AB_628423 | WB<br>1:500 |
| β-actin | Rabbit | Cell Signaling | 4967S<br>RRID: AB_330288 | WB<br>1:1000 |
|  | Mouse |  | 3700S<br>RRID: AB_2242334 | WB<br>1:1000 |
| Secondary antibodies |  |  |  |  |
| IgG DyLight 800 4x<br>PEG conjugate | Anti-rabbit | Cell Signaling | 5151S<br>RRID:<br>AB_10697505 | WB<br>1:1000 |
| IgG DyLight 680<br>conjugate | Anti-mouse |  | 5470S<br>RRID:<br>AB_10696895 | WB<br>1:1000 |
| Alexa Fluor 488 IgG | Anti-rabbit | Jackson ImmunoResearch<br>(West Grove, PA, USA) | 711-545-152<br>RRID: AB_2313584 | ICC<br>1:250 |
| Alexa Fluor 555 IgG | Anti-Mouse |  | 715-165-151<br>RRID: AB_2315777 | ICC<br>1:250 |

For biotinylation and western blotting, EZ-Link™ Sulfo-NHS-SS-Biotin (cat: 21331), NeutrAvidin Agarose Beads (cat: 29202), Pierce BCA Protein Assay Kit (cat: 23225), NuPAGE 4-12% Bis-Tris Mini Gels (cat: NP0323), NuPAGE MOPS SDS Running Buffer (cat: NP000102), and the XCell *SureLock* Mini-Cell system were purchased from ThermoFisher. The SpectraMax Plus 384 Absorbance Plate Reader was obtained from Molecular Devices Scientific (Waltham, MA, USA). PVDF membranes (cat: IPFL00005) were purchased from EMD Millipore. Intercept Blocking Buffer and LI-COR Odyssey XF were purchased from LI-COR Biosciences (Lincoln, NE, USA). VersaBlot total protein stain (cat: 33025/6-T) was purchased from Biotium (Fremont, CA).

For lactate dehydrogenase (LDH) assay, CyQUANT™ LDH Cytotoxicity Assay Kit (cat: C20300) was obtained from ThermoFisher. For co-immunoprecipitation (co-IP), Pierce Crosslink Magnetic IP/Co-IP Kit (cat: PI88805) was purchased from ThermoFisher. For immunocytochemistry, goat serum (cat: 005-000-121) was purchased from Jackson Immuno Research (West Grove, PA, USA), and Prolong Diamond Antifade Mountant solution with DAPI (cat: P36971) was purchased from ThermoFisher. For cell death analysis, SYTOX Green Nucleic Acid Stain (cat: S7020) was purchased from ThermoFisher. Imaging was performed using the LS720 LumaView microscope (Etaluma, Carlsbad, CA, USA).

### 2.3 Cell culturing

Primary glia cultures were prepared as previously described (Shimizu et al., 2011). In brief, postnatal 3-4-day old Sprague Dawley rat pups of both genders were rapidly decapitated, and brain cortices were isolated, combined, minced, and digested with 2.5% trypsin. Trypsin-containing dissection media was removed before tissue was then further digested with Deoxyribonuclease I (60 µg/ml, final concentration). Following several tissue titrations, cell suspension was centrifuged at 280 x g for 20 min. The supernatant was aspirated, and the pellet resuspended in Dulbecco’s modified Eagle’s (DMEM) media, supplemented with gentamicin (50 µg/ml) and 10% fetal bovine serum. After growth for 10 days *in vitro* (DIV), flasks were vigorously shaken to remove excess microglia, followed by trypsinization and centrifugation (280 x g for 20 min). Cell pellet was then resuspended in DMEM, counted on a Countess^TM^ III FL cell counter, and plated at 300,000 cells/well onto 6-well plates for western blotting; 2,000,000 cells/dish onto 100 mm dishes for immunoprecipitation; 600,000 cells/dish onto 60 mm dishes for internalization assays; 10,000 cells/well onto 96-well plates for glutamate uptake assays; and 30,000 cells/coverslip onto 12 mm coverslips for immunocytochemistry. All plates and dishes were coated with 0.2 mg/ml of poly-L-lysine prior to plating, except for 96-well plates, which were purchased pre-coated. Cultures were maintained for an additional 14 DIV before experiments.

For cell line cultures, COS-7 cells (ATCC, Manassas, VA, RRID: CVCL_0224) were maintained in DMEM containing 10% fetal bovine serum, 100 U/mL penicillin, and 100 μg/mL streptomycin in a humidified incubator at 37 °C with 5% CO_2_.

### 2.4 Cells transfections

COS-7 cells were transfected for immunoprecipitation and glutamate uptake assays using TransIT-LT1 transfection reagent. For immunoprecipitation, cells at ∼70% confluency were transiently transfected with 15 μg of plasmid DNA per dish (CMV-GLT-1-WT-3xFlag-T2A-EGFP and CMV-GLT-1-7KR-3xFlag-T2A-EGFP), plated at a density of 1,800,000 cells/dish and were lysed for immunoprecipitation two days later. For uptake assays, cells were transfected with 0.5 μg of plasmid DNA per well (empty vector CMV, to control for endogenous radiolabeled glutamate uptake, WT GLT-1 + empty vector, or GLT-1 + GLT-1 miR30 shRNA) and plated at a density of 50,000 cells/well. Uptake assays were performed two days later.

For glia cultures, siRNA transfections were performed using the DharmaFECT 3 transfection reagent. Transfections were performed 72 h before OGD with either Nedd4L (ID: M-092149-01-0010) or non-targeting control siRNA (ID: D-001206-13-05) purchased from Horizon.

### 2.5 Organotypic slice culturing

Organotypic slice cultures (OSCs) were prepared as described previously (Croft & Noble, 2018). In brief, brains were rapidly extracted from postnatal day 10-12 Sprague Dawley rat pups and placed into ice cold dissection medium. Tissue was sectioned into 200 μm slices using a McIlwain tissue chopper. Slices were cultured in 0.4 μm Millicell membrane inserts and maintained in culture medium consisting of Neurobasal A medium supplemented with 2% B27, 1% N2, 1% glutamine, 0.5% glucose, and 1% penicillin/streptomycin. Culture medium was replaced 24 h after slice preparation and subsequently refreshed by half-volume medium changes every other day thereafter for 10 days before experimentation.

### 2.6 Glia and OSC Lentiviral Transductions

To evaluate the role of ubiquitination in GLT-1 trafficking and ischemic recovery after OGD, glia cultures and OSCs were transduced with lentiviral constructs encoding GLT-1 C-terminal lysine residue mutants, which were obtained from VectorBuilder (Chicago, IL). Virus suspensions were aliquoted upon receival and stored at −80 °C until use. The following constructs were used: pLV-CMV-GLT-1-7KR-3xFlag-T2A-EGFP vector ID: VB240730-1621gjj, pLV-CMV-GLT-1-WT-3xFlag-T2A-EGFP vector ID: VB240812-1494exg, pLV-GFAP-GLT-1-miR30-GLT-1-7KR-3xFlag-T2A-mCherry vector ID: VB250904-1587nca, or pLV-GFAP-GLT-1-miR30-GLT-1-WT-T2A-3xFlag-mCherry vector ID: VB250904-1583gje and scramble virus pLV[miR30] EGFP/Puro EF1A mCherry:Scramble_miR30-shRNA vector ID: VB010000-9489dta. All virus preparations had titers ranging from 1 × 10^8^ to 1 × 10^9^ TU/mL.

2.6.1. Transduction of primary glial cultures: lentiviruses were added to the culture medium 7 days after secondary plating at a multiplicity of infection (MOI) of 10. Culture medium was completely replaced 24 h after transduction to remove residual virus.

2.6.2. Transduction of OSCs: 4–6 µL of lentivirus (MOI ∼8, depending on viral titer) was pipetted directly onto the surface of each hippocampal slice to allow dispersion across the entire slice. Culture medium was completely replaced 24 h after transduction.

### 2.7 Oxygen-Glucose Deprivation and Drug Treatments

Oxygen-glucose deprivation (OGD) insults were performed according to previous procedures (Goldberg & Choi, 1993). In brief, cell culture media was removed from the cells and replaced with degassed glucose-free DMEM. Plates and dishes were placed into the hypoxic chamber and de-oxygenated with a mixture of 10% H_2_/85% N_2_, and 5%CO_2_, then incubated at 37°C for 2 h (glial cultures) or 30 minutes (OSCs). Afterwards, the OGD media was removed and replaced with glucose-containing medium to allow reperfusion. For a subset of glial cultures, treatment with 80 μM Dynasore, 1 μM MG-132, 100 nM Bafilomycin, or 100 nM TAK-243 diluted into culture medium was performed following OGD. Cultures were returned to the incubator and maintained for 0-24 h of reperfusion. For a subset of OSC cultures, MK-801 at a final concentration of 10 μM was added to culture medium following OGD.

### 2.8 Glutamate uptake assays

Glutamate uptake assays were performed in glial cultures and COS-7 cells to measure the activity of glutamate transporters, as previously described (Fontana, 2018).

For glia, 96 well plates were washed using a plate washer, with PBS-CM (2.7 mM KCl; 1.2 mM KH_2_PO_4_, 138 mM NaCl; 8.1 mM Na_2_HPO_4_, added 0.1 mM CaCl_2_ and 1 mM MgCl_2_, pH 7.4). Uptake reactions were initiated by the addition of unlabeled L-glutamate and L-^3^H-glutamate (3.9-500 µM, final concentration, 99.96 % unlabeled and 0.04 % labeled). For uptakes assays performed in 24 well plates, cells were washed with PBS-CM and incubated with L-^3^H-glutamate (50 nM final concentration) or a mixture of unlabeled L-glutamate and L-^3^H-glutamate (250 µM, final concentration, 99.9 % unlabeled and 0.1 % labeled). Non-specific uptake was obtained in the presence of 150 µM TFB-TBOA. Incubation was carried out for 10 minutes at room temperature, then cells were lysed with 100 µL of EcoLite (+) scintillation cocktail and radioactivity was measured by scintillation counting.

For COS-7 cells, cultures were washed with PBS-CM and uptake reactions were initiated by addition of either L-³H-glutamate (50 nM final concentration) or a mixture of unlabeled L-glutamate and L-^3^H-glutamate (150 µM, final concentration, 99 % unlabeled and 1% labeled). After 10 min, uptake was terminated by two PBS-CM washes followed by addition of lysis buffer (1% sodium dodecyl sulfate / 0.1 M NaOH). Lysates were transferred to scintillation vials containing 3 mL scintillation cocktail, and radioactivity was measured by scintillation counting.

### 2.9 Lactate Dehydrogenase (LDH) Assay

Cell cytotoxicity following OGD was assessed using the CyQUANT™ LDH Cytotoxicity Assay Kit according to the manufacturer’s instructions. LDH reagent was added directly to culture wells to measure LDH released into the medium. As a positive control, lysis buffer supplied with the kit was applied to selected wells to induce maximal cell death. This allowed calculation of percent cytotoxicity relative to maximum LDH release for each experimental condition.

### 2.10 Sodium-Potassium ATPase (NKA) Assay

Activity levels of Sodium Potassium ATPase (NKA) were measured using a Fluorimetic SensoLyte FDD Protein Phosphatase Assay Kit (Anaspec, Japan), according to manufacturer’s instructions. A fluorescence microplate reader was used to detect fluorescein emission at 528 nm with excitation at 485 nm. Fluorescein emissions were recorded every 5 minutes after reaction started, for a total of 30 minutes. Ouabain, a NKA inhibitor (Tocris, Bristol, United Kingdom) was added to select wells at a final concentration of 15 mM to obtain the background. Enzymatic activity was determined by calculating the difference between fluorescein signal measured with and without the presence of ouabain.

### 2.11 Cell Surface Biotinylation

Surface biotinylation procedures were performed in glial cultures and OSCs to quantify changes in GLT-1 surface protein expression after OGD insults. Cultures were washed and treated with 1 mg/ml of EZ-Link™ Sulfo-NHS-SS-Biotin (ThermoScientific), diluted in PBS-CM for glial cultures or artificial cerebrospinal fluid (aCSF: 125 mM NaCl, 2.5 mM KCl, 1.2 mM NaH_2_PO_4_, 1.2 mM MgCl_2_, 2.4 mM CaCl_2_, 26 mM NaHCO_3_, and 11 mM glucose) for OSCs. Reactions proceeded for 30 minutes on ice and quenched with 100 mM glycine. Glial cultures were lysed with 500 µl TNE-lysis buffer (10 mM Tris-HCL, 1 mM EDTA, 150 mM NaCl, and 1% Triton) and incubated on a rocker at 4°C for 30 minutes. OSCs tissue was collected and resuspended in ice cold RIPA buffer (10 mM Tris, pH 7.4, 150 mM NaCl, 1.0 mM EDTA, 1% Triton-X-100, 0.1% SDS, 1% Na deoxycholate) to break up tissue and incubated on a rocker at 4°C for 30 minutes. Lysates were collected and centrifuged for 10 min at 16,000 x g at 4°C. A portion of the lysate (40 µl, ∼7% of total volume) was set aside for protein quantification and 32 µl was reserved for total lysate analysis. The remaining lysate (400 µl) was incubated overnight with 50 µl of NeutrAvidin Agarose Resin to isolate biotinylated protein. Bead-containing lysates were then centrifuged at 4°C and washed 3 times with TNE lysis buffer for glia cultures (or RIPA buffer for OSCs) and once with PBS-CM before being prepared for western blot analysis.

### 2.12 Western blot approaches

Protein content of lysates was determined using a Pierce BCA Protein Assay Kit. Samples were prepared by the addition of NuPage LDS sample buffer and DTT and heated at 65°C. Equal amounts of protein (3–4 μg) were loaded into NuPAGE 4-12% Bis-Tris Mini Gels, and electrophoresis was run for 60 mins at a constant 200 V using NuPAGE MOPS SDS Running Buffer with the XCell *SureLock* Mini-Cell system. Proteins were transferred onto PVDF membranes using a Bio Rad TransBlot Turbo Transfer system, according to manufacturing protocol. Membranes were blocked with Intercept Blocking Buffer for 1 h, followed by overnight incubation with primary antibodies (listed in Table 1) at 4°C. For glial cultures, monoclonal β-actin (mouse or rabbit, matched to the primary antibody species) was used as a loading control (Table 1). For OSCs, total protein loading was assessed using VersaBlot™ total protein stain. The following day, primary antibodies were removed, and membranes were washed with PBS-Tween (PBS-T) and incubated with fluorescent-conjugated secondary antibodies for 60 min at room temperature. After additional PBS-T washes, membranes were imaged using a LI-COR Odyssey XF imaging system.

### 2.13 Co-Immunoprecipitation

To determine protein interactions between GLT-1 and post-translational modifiers after OGD, co-IP was performed using Pierce Crosslink Magnetic IP/Co-IP Kit according to manufacturer’s protocol. In brief, 4-24 h following OGD, glial cultures were washed and lysed with IP/lysis buffer. Samples were centrifuged at 16,000 x g, and supernatants were normalized to equal protein concentrations. Magnetic beads were crosslinked to antibody (GLT-1: 0.04 µg/µl, IgG: 0.02 µg/µl, Flag: 0.01 µg/µl) via incubation with 20 µM disuccinimidyl suberate. 500 µl of lysate was added to crosslinked beads and incubated at room temperature for 1 h. The remaining lysate was set aside for analysis of input. Beads were washed with IP/lysis buffer, and protein was eluted with 60 µl of elution buffer prior to western blot analysis (as described above).

### 2.14 Immunocytochemistry

To determine colocalization of GLT-1 with EEA1 following OGD or to characterize cell type composition and glutamate transporter distribution in cultures, coverslips were washed and fixed with 4% PFA and permeabilized with 0.1% Triton-X in PBS. Samples were blocked for 30 min with 8% goat serum, followed by overnight incubation at 4°C with primary antibodies diluted in blocking solution. The next day, primary antibodies were removed, coverslips were washed with PBS and then incubated with secondary antibodies (anti-rabbit Alexa-488 and anti-mouse Alexa 555), diluted in blocking buffer for 45 min in the dark at room temperature. Coverslips were washed with PBS, rinsed in Milli Q water to remove any excess salt on the coverslips, and mounted on microscope slides using 7 μl of Prolong Diamond antifade mountant with DAPI. Coverslips were imaged using the Olympus FluoView FV3000 at a 100X objective. Five fields were captured for duplicate coverslip of each condition across four independent experiments.

### 2.15 SYTOX green staining and cell death analysis

To assess the neuroprotective potential of inhibiting GLT-1 C-terminal PTMs during OGD, cell death in hippocampal OSCs was quantified using SYTOX Green staining. SYTOX Green was added to the OSC culture medium at a final concentration of 100 µM and incubated for 30 min at 37 °C. Prior to OGD, baseline images were acquired using 488 nm (green) and 555 nm (red) excitation channels via a LS720 Lumaview microscope. Following OGD, slice cultures were transferred to Sytox Green–containing wells and imaged hourly for up to 24 h post-insult.

### 2.16 Data and statistical analysis

All data analysis was performed using GraphPad Prism 10.4 (GraphPad Software, La Jolla, CA, USA).

For kinetic analysis, specific uptake values were obtained by subtracting background values (in presence of TFB-TBOA), converted to nmol/µg/min, and analyzed assuming Michaelis−Menten kinetics for calculation of V_max_ and K_m_. K_m_ values were log-transformed before statistical comparisons.

Quantification of Western blot band intensities of both surface (biotinylated) and total EAAT2 monomers (∼65 kDa) was performed using LI-COR Image Studio software. Protein band intensities were normalized to actin as a loading control (glia) or to total protein stain (OSCs). For surface-expressed biotinylated proteins, values were expressed as a percentage of total protein and normalized within each blot.

Immunocytochemistry images were analyzed in FIJI (ImageJ). Colocalization between fluorescence channels was assessed using the JaCoP (Just Another Colocalization Plugin) by calculating the intensity correlation quotient (ICQ) for each image. ΔICQ values were obtained by normalizing individual ICQ measurements to the average control ICQ within each biological replicate, and mean ΔICQ values were plotted for each condition.

For SYTOX Green cytotoxicity analysis, pre-OGD and post-OGD (24 h reperfusion) images of hippocampal OSCs were analyzed using FIJI (ImageJ). The same three regions of interest (ROIs) within each hippocampal region were selected in the corresponding pre- and post-OGD images. Integrated density was calculated by averaging triplicate ROIs after background subtraction. Data were expressed as a percentage increase relative to the corresponding pre-OGD image.

For immunocytochemistry and cytotoxicity studies, the experimenter was blinded during data analysis.

Before parametric statistical analyses were performed, data normality was assessed using the Shapiro–Wilk test and inspection of QQ plots. Datasets were also screened for outliers using the ROUT method (Q = 1%), and identified outliers were excluded prior to statistical analysis. Inclusion or exclusion of outliers did not alter the overall statistical conclusions. Depending on the experimental design, statistical comparisons were performed using a two-tailed unpaired Student’s *t* test, one-way ANOVA, or two-way ANOVA, followed by Dunnett’s multiple comparisons test or Fisher’s least significant difference (LSD) test for planned comparisons, as indicated in the figure legends. A summary of statistical analyses for each figure is provided in Table 2.

**Table 2:** Statistical analysis of the main figures of this study.

| Figure |  | Statistical Test | Adjusted P | F-value: F (DFn, DFd) |
| --- | --- | --- | --- | --- |
| 1 | E | Mixed-effects model<br>(Dunnett's *post-hoc*) | 0.0004 (0 h)<br>0.0053 (4 h)<br>0.0006 (24 h) | Reperfusion effect: F (3,9) = 31.14<br>$p < 0.0001$ |
| | G | | 0.0103 (4 h)<br>0.0003 (6 h)<br>0.0428 (24 h) | Reperfusion effect: F (6,17) = 9.078<br>$p = 0.0002$ |
| 2 | B | Two-tailed unpaired *t*-test | 0.0038 | n/a |
|  | E |  | 0.0089 | n/a |
| | F | 2-Way ANOVA | 0.0020 (vehicle vs DYN,<br>OGD group)<br>0.0004 (control vs OGD,<br>in vehicles) | OGD Effect: F (1, 6) = 15.13,<br>$p = 0.0081$<br>Treatment Effect: F (1, 6) = 16.27<br>$p = 0.0069$<br>OGD vs. Treatment effect: F (1, 6) = 11.2<br>$p = 0.0154$ |
|  | I | Two-tailed unpaired *t* test | 0.0473 | n/a |
| 3 | B | 2-Way ANOVA<br>(Fisher's LSD *post-hoc*) | 0.0029 (control vs OGD,<br>in vehicles) | OGD Effect: F (1, 6) = 8.418<br>$p = 0.0273$ |
| | C | | 0.0037 (vehicle vs MG-132, control group) | Treatment Effect: F (1, 6) = 20.91<br>$p = 0.0038$ |
| | D | | 0.0002 (control vs OGD,<br>in vehicles) | OGD Effect: F (1, 6) = 39.44<br>$p = 0.0008$ |
| | F | | 0.0002 (control vs OGD,<br>in vehicles)<br>0.0052 (vehicle vs Baf +<br>MG-132, OGD group) | OGD Effect: F (1, 6) = 11.60<br>$p = 0.0144$<br>OGD vs. Treatment effect: F (1, 6) =<br>21.47<br>$p = 0.0036$ |
| | G | | 0.0095 (vehicle vs Baf +<br>MG-132, OGD group) | Treatment Effect: F (1, 6) = 35.30<br>$p = 0.0010$ |
| 4 | B | Two-tailed unpaired *t* test | 0.0096 | n/a |
|  | C |  | <0.0001 |  |
| 5 | B | 2-Way ANOVA<br>(Fisher's LSD *post-hoc*) | 0.0311 (control vs OGD,<br>in WT group)<br>0.0491 (WT vs 7KR, in<br>OGD group) | OGD x Treatment effect: F (1, 6) = 7.357<br>$p = 0.0350$ |
|  | D | Mixed-effects model<br>(Fisher's LSD *post-hoc*) | 0.0096 (control vs OGD,<br>in WT group)<br>0.0172 (WT vs 7KR, in<br>OGD group) | OGD x Treatment Effect: F (1, 7) = 5.751 |
| 6 | B | 2-Way ANOVA<br>(Fisher's LSD *post-hoc*) | 0.0046 (control vs OGD,<br>in WT group)<br>0.0027 (WT vs 7KR, in<br>OGD group) | OGD Effect: F (1, 6) = 9.463<br>$p = 0.0218$<br>OGD vs. Treatment effect: F (1, 6) =<br>9.799<br>$p = 0.0203$ |
| | D | | 0.0051 (control vs OGD,<br>in WT group)<br>0.0016 (WT vs 7KR, in<br>OGD group) | OGD Effect: F (1, 6) = 20.99<br>$p = 0.0038$<br>Treatment Effect: F (1, 6) = 17.24<br>$p = 0.0060$ |
| | F | | 0.0021 (control vs OGD,<br>in WT group)<br>0.0128 (WT vs 7KR, in<br>OGD group) | OGD Effect: F (1, 6) = 27.84<br>$p = 0.0019$<br>Treatment Effect: F (1, 6) = 4.667<br>$p = 0.0741$<br>OGD vs. Treatment effect F (1, 6) = 3.986<br>$p = 0.0929$ |
| 7 | B | 2-Way ANOVA<br>(Fisher's LSD *post-hoc*) | 0.0012 (control vs OGD,<br>in vehicles)<br>0.0076 (vehicle vs TAK-243, OGD group) | OGD Effect: F (1, 6) = 7.603<br>$p = 0.0330$<br>OGD vs. Treatment effect: F (1, 6) =<br>14.85<br>$p = 0.0084$ |
| | D | | 0.0067 (control vs OGD,<br>in vehicles) | OGD Effect: F (1, 6) = 5.908<br>$p = 0.0511$ |
| | | | | OGD vs. Treatment effect: $F(1, 6) = 4.863$<br>$p = 0.0696$ |
| 8 | B | Two-tailed unpaired $t$ test | 0.0044 (control vs OGD) | n/a |
| | C | Two-tailed unpaired $t$ test | 0.0462 (control vs OGD) | n/a |
| | G | One-Way ANOVA<br>(Dunnett's multiple comparisons) | CA1: 0.0235 (WT vs 7KR)<br>0.0003 (Scramble MK-801 vs WT)<br>CA3: 0.0383 (Scramble MK-801 vs WT) | CA1: $F(2, 11) = 15.70$<br>$p = 0.0006$<br>CA3: $F(2, 11) = 4.314$<br>$p = 0.0414$ |

## 3 Results

### 3.1 Glutamate Uptake Velocity and GLT-1 Surface Expression are Decreased following Oxygen-Glucose Deprivation in Primary Glia Cultures

We examined the effects of ischemic insult followed by varying durations of reperfusion on primary glial cultures. Characterization by immunocytochemistry revealed that cultures consisted of 90-95% glia (approximately 70% astrocytes and 30% microglia) and 5-10% neurons, that had robust expression of both GLAST and GLT-1 (Supplemental Figure 1A). Importantly, the presence of a subset of neurons in our culture system is necessary, as expression of GLT-1 is dependent on neuronal-secreted factors (Gegelashvili et al., 1997; Schlag et al., 1998; Zelenaia et al., 2000). Additionally, the presence of microglia in these cultures permits microglia–astrocyte interactions that may influence astrocytic responses to injury (Yang et al., 2025). While microglial signaling has been reported to influence astrocytic transporter regulation, the astrocyte-enriched nature of our cultures suggests that the observed changes in GLT-1 trafficking largely reflect astrocytic responses to ischemic stress. Further pharmacological characterization using selective glutamate transporter inhibitors, at doses selective for inhibiting GLT-1 (1 µM WAY 21363), GLAST (10 µM UCPH-101) or all of the transporters (100 µM TFB-TBOA), indicated that glutamate uptake in these cultures is primarily mediated by GLT-1 (∼80%), with a smaller contribution from GLAST (∼20%) (Supplemental Figure 1B). A small fraction of glutamate uptake remained following treatment with 100 μM TFB-TBOA, suggesting possible presence of Na⁺-independent transporters, such as the system x_c_ ^-^ cystine/glutamate antiporter, which has been reported in astrocytes (Bridges et al., 2012). These results confirm that our cultures are astrocyte-enriched mixed glial cultures exhibiting robust GLT-1 expression, which we refer to as glial cultures for the remainder of the study.

We next examined the effects of ischemic insult on GLT-1 regulation. Primary glia cultures were subjected to oxygen-glucose deprivation (OGD) followed by varying lengths of reperfusion, after which glutamate uptake and GLT-1 surface expression were analyzed (Figure 1A). We first wanted to determine the effect of varying OGD insult lengths on transporter expression and activity. OGD insults of 1 and 2 h resulted in the most robust decreases in V_max_ and surface GLT-1 at 24 h reperfusion, while total GLT-1 levels remained unchanged. On the other hand, a 30 min OGD insult reduced V_max_ without altering GLT-1 surface expression (Supplemental Figure 2A-2C). Expression of GLAST, the other primary astrocytic-expressed glutamate transporter subtype, was unaffected by OGD (Supplemental Figure 2D and 2E). Based on these results, 2 h OGD was used for all subsequent experiments.

**Figure 1.**
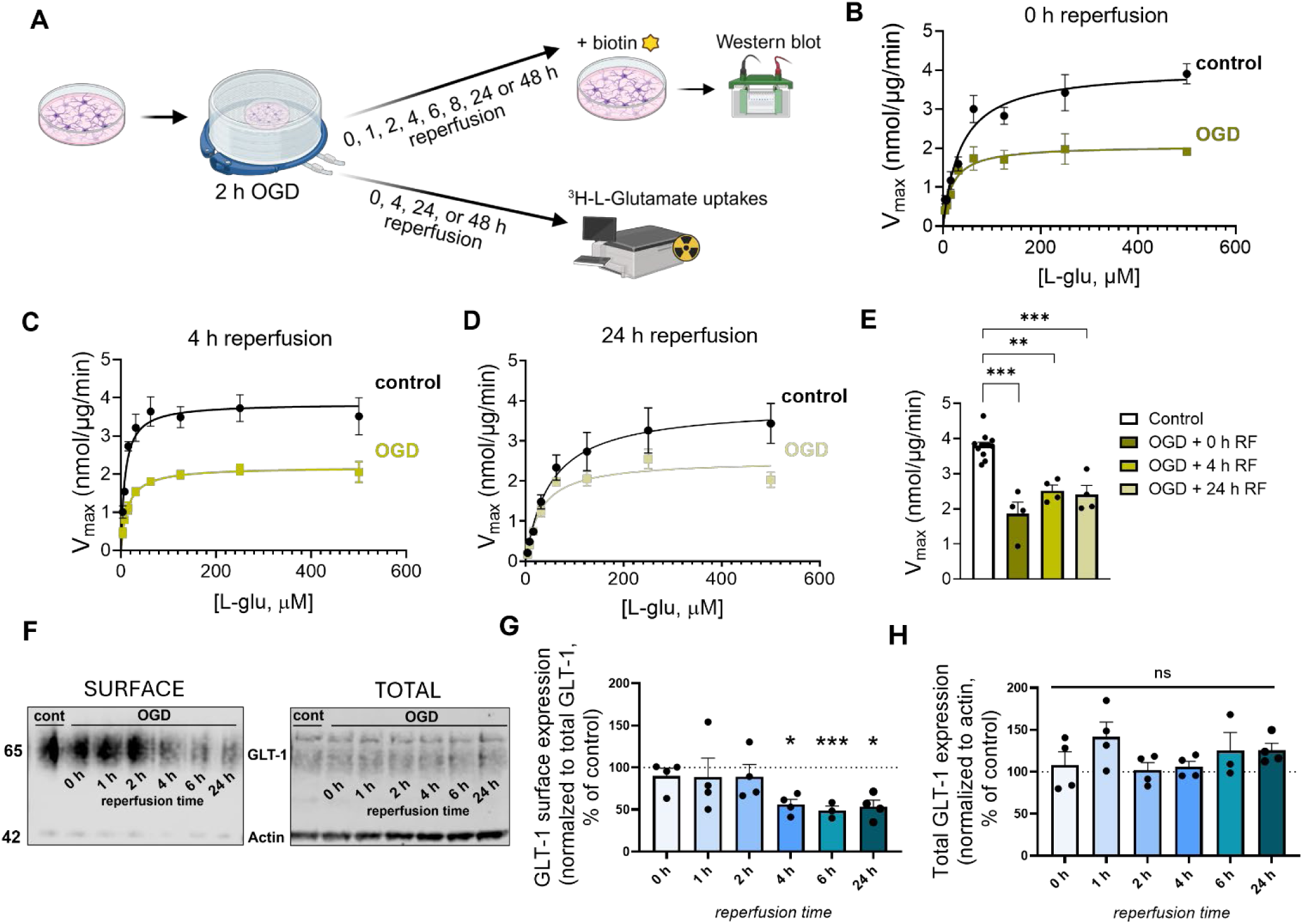
Oxygen-glucose deprivation reduces glutamate uptake velocity and GLT-1 surface expression in primary glia cultures. **A.** Schematic of experimental paradigm. Glial cultures were subjected to 2 h oxygen-glucose deprivation (OGD), or maintained as non-insult (control), followed by reperfusion for the indicated time points (0-24 h). Glutamate uptake and GLT-1 surface expression were subsequently assessed. **B-D.** Representative Michaelis-Menten saturation curves of L-^3^H-glutamate uptake measured at 0 h, 4 h or 24 h of reperfusion, respectively. **E**. Quantification of V_max_ values normalized to nmol/μg/min (n=4-12 independent cell culture preparations per group with 11 replicates averaged) **F**. Representative immunoblots of surface expression (left) and totals (right) of GLT-1 (∼62 kDa) at 0 h, 1 h, 2 h, 4 h, 6 h, and 24 h reperfusion following OGD insult or non-insult controls. **G-H**. Quantification of surface (**G**) and total (**H**) GLT-1 expression across varying reperfusion time points (n=4-7 independent cell culture preparations per group with technical duplicates averaged). Data in panels B-E and G-H are presented as mean ± SEM. Statistical analysis of data in panels E, G and H were performed using a mixed-effects model (REML), with experimental condition treated as a fixed effect and experimental day as a random effect to account for pairing within experiments. *Post hoc* comparisons versus non-insult controls were performed using Dunnett’s multiple-comparisons test to control the family-wise error rate., * *p* < 0.05, ** *p* < 0.01 and *** *p* < 0.001, ns=non-significant.

To have a better temporal understanding of how GLT-1 is affected in the earlier stages of reperfusion, glutamate transport kinetics were measured immediately after (0 h reperfusion), and at 4 or 24 h reperfusion using 3.9-500 µM L-^3^H-glutamate. Representative Michaelis–Menten saturation curves for each time point are shown in Figure 1B-D. OGD significantly reduced V_max_ by approximately 39%, 30% and 32% at 0, 4, and 24 h reperfusion respectively, compared to non-insult controls (Figure 1E, *p <* 0.001 for 0 and 24 h, *p <* 0.01 for 4 h). No significant changes were observed on K_m_ (mixed-effects model, (F (3,9) = 1.676, *p* = 0.2407). Importantly, these changes occurred without effects on cell cytotoxicity (Supplemental Figure 3A).

Next, to determine whether reduced uptake was associated with altered transporter trafficking, we measured surface expression levels of GLT-1 at the same reperfusion time points. Surface protein biotinylation specificity was confirmed by the absence of intracellular protein signal, β-actin, in the biotinylated fraction (Figure 1F). Surface GLT-1 levels were significantly downregulated by approximately 44% at 4 h reperfusion and remained decreased at 24 h (Figure 1F and 1G, *p* < 0.001 for 6 h reperfusion and *p* < 0.05 for 4 h and 24 h reperfusion), whereas total GLT-1 expression levels remained unchanged (Figure 1F and 1H).

The impaired glutamate uptake observed immediately after ischemic insult (0 h reperfusion) is likely attributable to reduction of ATP stores that are necessary for secondary active transport of glutamate. This may explain the significant reduction in V_max_ at 0 h reperfusion despite unchanged surface GLT-1 levels. To test this, we measured enzymatic activity of the sodium-potassium ATPase (NKA), the primary ATPase that maintains the ionic gradients necessary for glutamate transport. NKA activity was reduced by approximately 42% at 0 h reperfusion, but recovered at later timepoints (Supplemental Figure 3B, *p* < 0.01). This transient decrease in NKA activity likely contributes to the reduction in V_max_ observed immediately after OGD, despite unchanged GLT-1 surface expression levels. Both GLT-1 surface expression levels (Supplemental Figure 4A–C) and glutamate uptake velocity (Supplemental Figure 4D and 4E) returned to baseline by 48 h reperfusion. Together, these findings indicate that impaired glutamate clearance observed during mid to late-stage reperfusion is primarily driven by a reduction in GLT-1 surface availability.

### 3.2 GLT-1 is Internalized through the Early Endosomal Pathway following OGD

To determine whether GLT-1 internalization contributes to its downregulation following OGD, we performed an endocytic biotinylation assay. Surface proteins were biotinylated prior to OGD insult to enable tracking throughout ischemic reperfusion, and internalized GLT-1 was measured following removal of remaining biotin-labeled surface proteins using a glutathione buffer (Figure 2A). First, we validated the assay by treating cells for 90 min with 100 nM of phorbol-12-myristate 13-acetate (PMA), a well-established inducer of GLT-1 internalization via activation of protein kinase C (Susarla & Robinson, 2008). Upon treatment with PMA and glutathione reduction, we observed a decrease in GLT-1 surface expression levels (Supplemental Figure 5A) that coincided with an increase in internalized GLT-1 (Supplemental Figure 5B, *p* < 0.05), confirming the sensitivity of the assay. Next, we performed this experiment in our model and focused on the 4 h reperfusion timepoint, when GLT-1 surface was first found to be downregulated (Figure 1G), and the 24 h time point. We observed an increase in internalized GLT-1 following 4 h reperfusion (Figure 2B, *p* < 0.05), but not at 24 h (Figure 2C). On the other hand, the total amount of pre-biotinylated surface GLT-1 remained unchanged at 4 h (Figure 2D), suggesting that degradation of internalized pools had not occurred yet. In contrast, pre-labeled GLT-1 was significantly reduced at 24 h reperfusion, suggesting degradation of the internalized pools (Figure 2E). To evaluate whether preventing GLT-1 internalization restores transporter surface expression, cultures were treated with Dynasore immediately following OGD. Dynasore was used at 80 µM, a concentration previously shown to inhibit dynamin-dependent endocytosis in mammalian cells without cytotoxicity (Macia et al., 2006; Kirchhausen et al., 2008). Dynasore treatment prevented the OGD-induced reduction in surface GLT-1, compared to vehicle-treated controls, while total GLT-1 expression remained unchanged (Figure 2F and 2G). Consistent with this findings, immunofluorescence staining and confocal imaging demonstrated increased colocalization of GLT-1 with the early endosome marker EEA1 at 4 h reperfusion following OGD (Figure 2H and 2I), but not 24 h (Figure 2J). Together, this data suggests that OGD triggers dynamin-dependent internalization of GLT-1 into early endosomal compartments, which precedes degradation and contributes to a reduced transporter surface expression following OGD.

**Figure 2.**
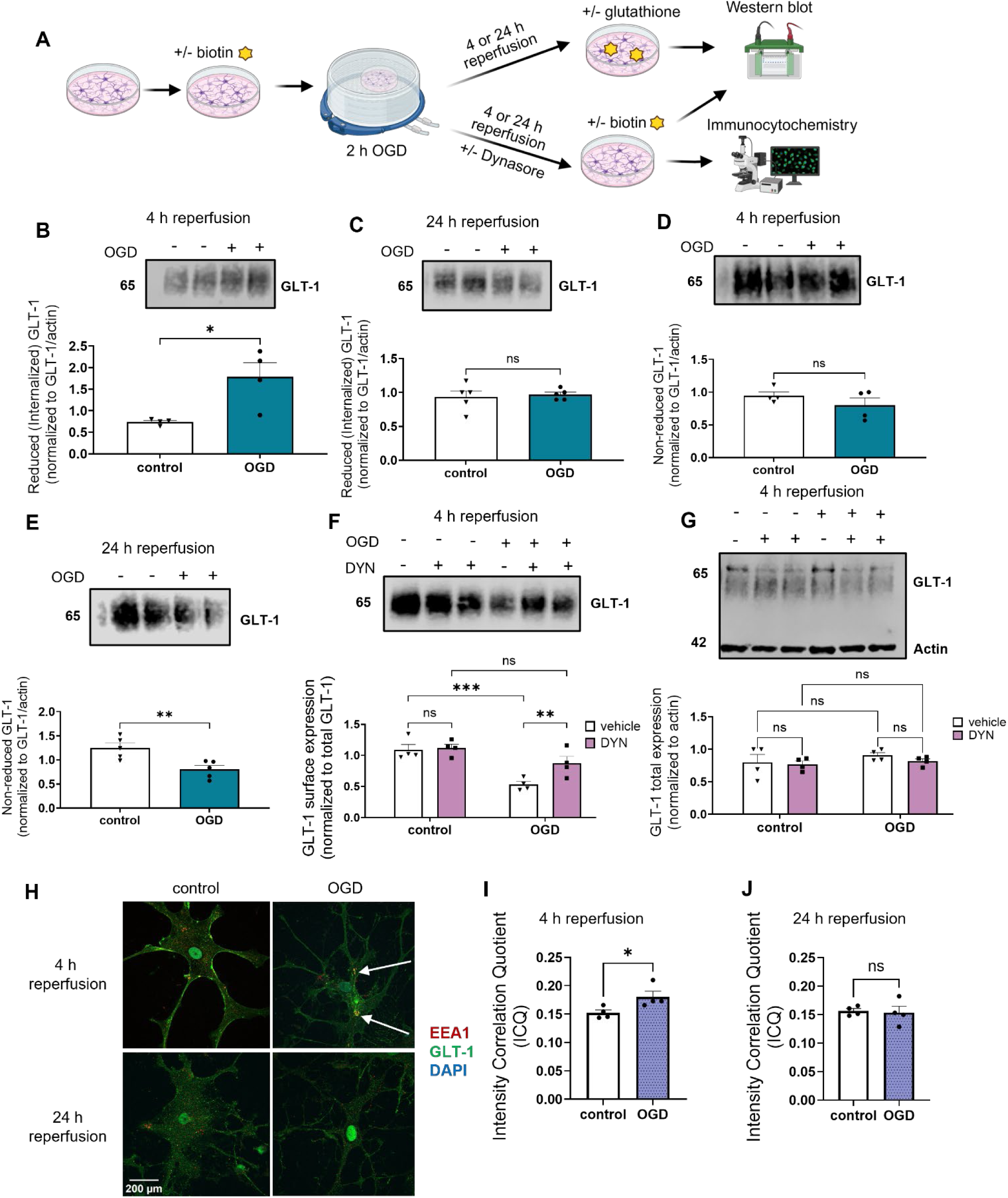
OGD induces GLT-1 internalization through the early endosomal pathway. **A.** Schematic of experimental paradigm. For endocytic biotinylation, assays, a subset of cultures was biotinylated prior to OGD, followed by reduction with glutathione and immunoblotting at the indicated reperfusion times. Another subset of cultures was treated with Dynasore (DYN) immediately post-OGD, followed by biotinylation and immunoblotting. Additional OGD cultures without further treatment were used for immunocytochemistry. **B and D.** Representative immunoblots (top) and protein quantification (bottom) of internalized, pre-labeled GLT-1 measured by endocytic biotinylation at 4 h (**B**) or 24 h reperfusion (**C**). **D and E**. Representative immunoblot (top) and protein quantification (bottom) of total pre-labeled GLT-1 (remaining surface and internalized pools) at 4 h (**D**) or 24 h reperfusion (**E**) (n=4-5 independent cell culture preparations per group; technical duplicates averaged). **F-G**. Representative immunoblot (top), and protein quantification (bottom) of surface (**F**) and total (**G**) GLT-1 following 4 h reperfusion with Dynasore (DYN, 80 µM) applied immediately upon termination of OGD and maintained throughout the reperfusion period (n=4 independent cell culture preparations per group; technical duplicates averaged). **H.** Representative immunofluorescent images showing GLT-1 colocalization with the early endosome marker EEA1 at 4 or 24 h reperfusion. Scale bar: 200 µm. **I-J**. Intensity Correlation Quotient (IQC) analysis quantifying GLT-1 and EEA1 colocalization at 4 h (**I**) and 24h (**J**) reperfusion (n=4 independent cell culture preparations per group of duplicates; 5 cells image per coverslip). Data are presented as mean ± SEM. Statistical analyses were performed using two-tailed unpaired *t* tests (B-E, I-J), and Two Way ANOVA followed by Fisher’s LSD *post-hoc* test (F-G). * *p* < 0.05, ** *p* < 0.01, *** *p* < 0.001.

### 3.3 GLT-1 is Degraded through Multiple Degradation Pathways following OGD

To determine the post-endocytic processing of internalized GLT-1 following OGD, cultures were treated with proteasomal or lysosomal pathway inhibitors (MG-132 and Bafilomycin, respectively), after 2 h OGD (Figure 3A). MG-132 (1 μM) and Bafilomycin A1 (100 nM) were used at concentrations cited in prior studies examining the regulation of glutamate transporters (Zhang et al., 2017; Iovino et al., 2022). MG-132 does not affect GLT-1 surface expression by itself, but prevents the OGD-induced downregulation of GLT-1 surface expression upon proteasome inhibition (Figure 3B). MG-132 treatment also increased total GLT-1 levels by approximately twofold, suggesting that proteasomal degradation contributes to basal GLT-1 turnover (Figure 3C, *p* < 0.01 vehicle vs MG-132 in control group). Similar results were observed with the lysosomal inhibitor Bafilomycin A1 (Baf), which also failed to prevent the OGD-induced decrease in surface GLT-1 (Figure 3D). In contrast to MG-132, Baf treatment had no effect on total GLT-1 expression levels (Figure 3E). Interestingly, co-treatment with both proteasomal and lysosomal inhibitors fully prevented OGD-induced downregulation of surface GLT-1 (Figure 3F, *p* < 0.01, vehicle vs Baf + MG in OGD group), and significantly increased total GLT-1 levels (Figure 3G). Together, these results suggest that internalized GLT-1 is degraded through both proteasomal and lysosomal pathways following OGD.

**Figure 3.**
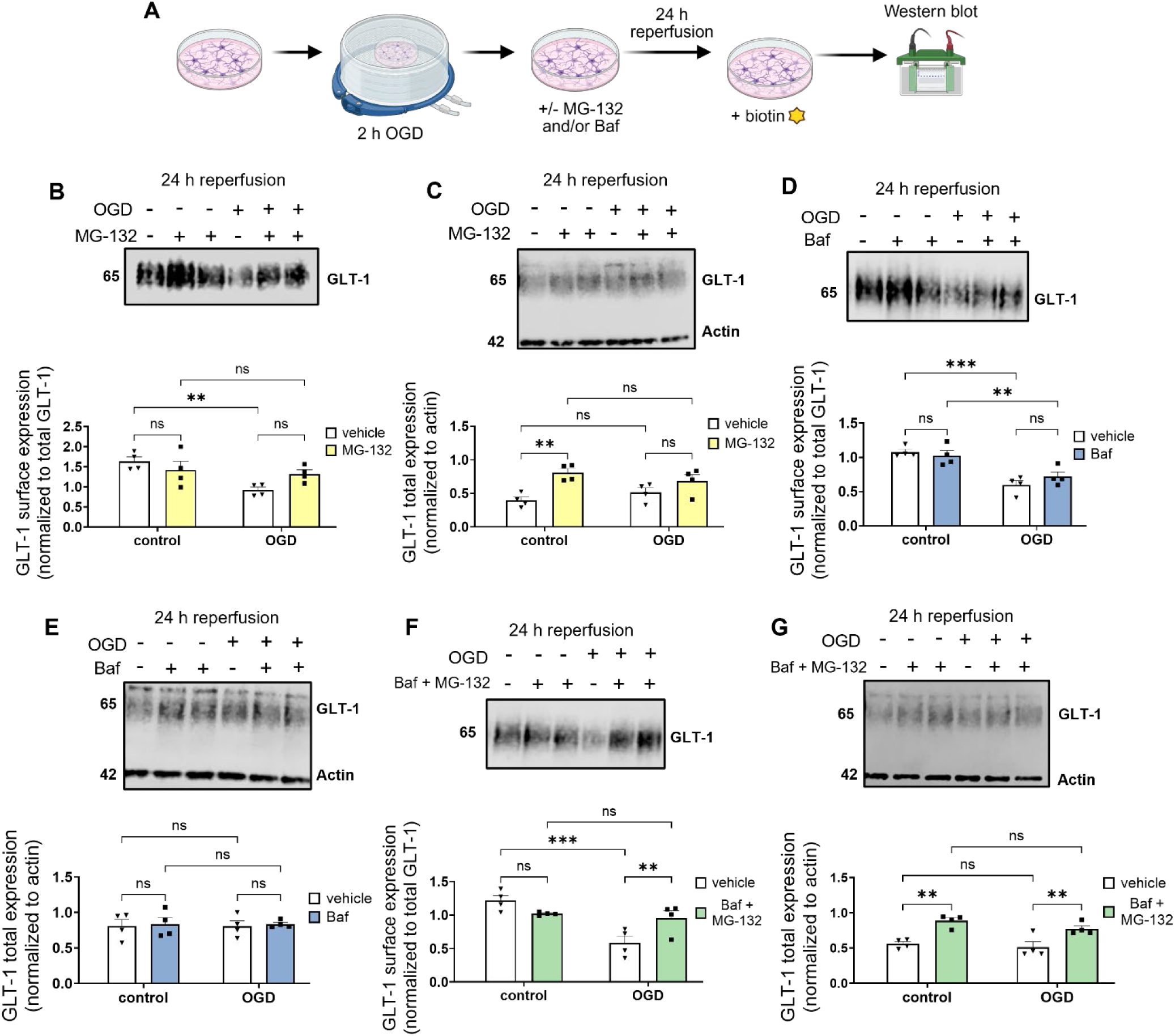
Internalized GLT-1 is degraded through lysosomal and proteasomal pathways. **A.** Schematic of experimental paradigm. MG-132 or Bafilomycin (Baf) were applied immediately after OGD and maintained throughout reperfusion, followed by surface biotinylation and immunoblotting at the indicated time points. **B-C.** Representative immunoblots (top), and protein quantification (bottom) of surface (**B**) and total (**C**) GLT-1 following 24 h reperfusion with 1 µM MG-132. **D-E.** Representative immunoblots (top), and protein quantification (bottom) of surface (**D**) and total (**E**) GLT-1 following 24 h reperfusion with 100 nM Baf applied immediately upon termination of OGD. **F-G.** Representative immunoblots (top), and protein quantification (bottom) of surface (**F)** and total (**G**) GLT-1 following 24 h reperfusion with combined 100 nM Baf and 1 µM MG-132 treatment. Data are presented as mean ± SEM (n=4 independent cell culture preparations per group; technical duplicates averaged). Statistical analyses were performed using Two Way ANOVA followed by Fisher’s LSD *post-hoc* test. * *p* < 0.05, ** *p* < 0.01, *** *p* < 0.001.

### 3.4 Ubiquitination and SUMOylation of GLT-1 Increase following OGD

Previous reports have identified several post-translational modification, notably at C-terminal lysine residues, as regulators of GLT-1 trafficking (Sheldon et al., 2008; Gonzalez-Gonzalez et al., 2008; Martínez-Villarreal et al., 2012; Foran et al., 2014). Therefore, to investigate whether ischemic insult alters post-translational modifications of GLT-1, we performed co-immunoprecipitation of GLT-1 followed by immunoblotting for ubiquitin and SUMO-1, which both occur at lysine residues of target protein. Following OGD, cell lysates were collected at the indicated reperfusion time points and GLT-1 was immunoprecipitated with eluants immunoblotted for ubiquitin and SUMO-1. At 4 h reperfusion, no change in GLT-1 SUMOylation was detected (Figure 4A), whereas a significant increase was observed at 24 h reperfusion (Figure 4B). Consistent with literature, SUMO-1 lysates appeared as a single band at 17 kDA, representing the pool of unconjugated SUMO-1 as conjugated forms were most likely undetectable without enrichment (Park-Sarge & Sarge, 2010). Conversely, GLT-1 ubiquitination was significantly increased at 4 h reperfusion (Figure 4C) but returned to baseline levels by 24 h reperfusion (Figure 4D). As GLT-1 internalization was first observed at 4 h reperfusion, the early increase in ubiquitination supports ubiquitin-dependent modification as a potential driver of early transporter downregulation and degradation of GLT-1 following OGD, while not excluding a potential role for SUMOylation at later stages of transporter regulation.

**Figure 4.**
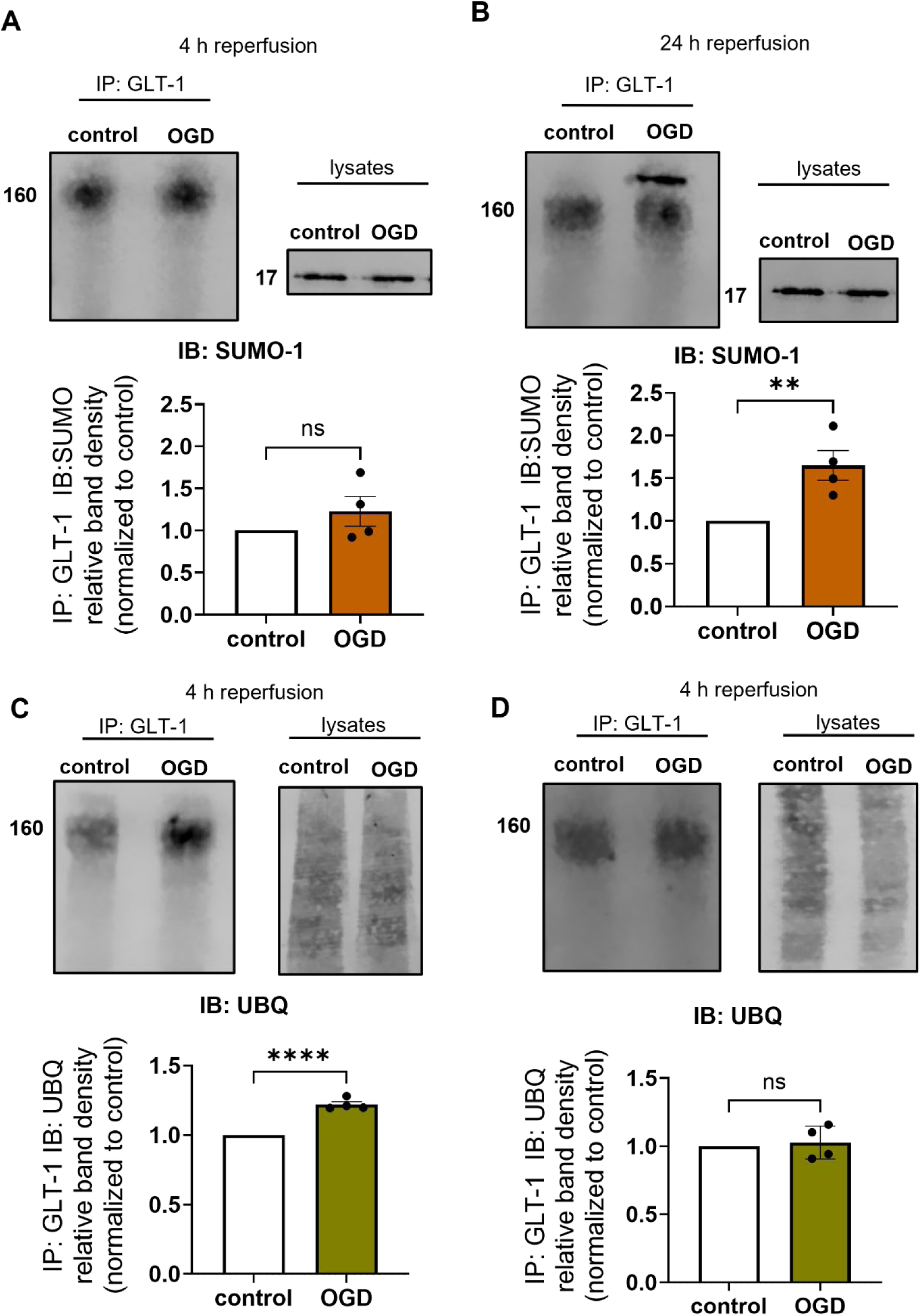
OGD induces lysine-directed post-translational modification of GLT-1. Following OGD, GLT-1 was immunoprecipitated from glial lysates and immunoblotted for ubiquitin (P4D1) or SUMO-1. **A-B**. Representative immunoblot with IP and total lysate input (top) and protein quantification (bottom) of SUMOylated GLT-1 at 4 h (**A**) or 24 h (**B**) reperfusion. **C-D**. Representative immunoblots (top) and protein quantification (bottom) of ubiquitinated GLT-1 following 4 h (**C**) or 24 h (**D**) reperfusion (n=4 independent cell culture preparations per group; technical duplicates averaged). Data is presented as fold change relative to non-insult control ± SEM. Statistical analyses were performed using two-tailed unpaired *t* test. ** *p* < 0.01, **** *p* < 0.0001, ns= non-significant.

### 3.5 Inhibition of C-terminal Lysine PTMs Prevent GLT-1 Internalization and Restore Functional Uptake

To determine whether lysine-directed modifications mediate GLT-1 downregulation following OGD, we generated a lentiviral construct encoding a C-terminal lysine-to-arginine mutant (GLT-1 7KR). To enable probing specifically for virally expressed GLT-1 constructs, an N-terminal Flag tag was added into both the 7KR mutant (Flag-GLT1 7KR) and wild-type GLT-1 (Flag–GLT-1 WT) (Figure 5A). First, to verify that the 7KR construct effectively impeded GLT-1 ubiquitination, COS-7 cells were transfected with Flag-GLT-1 7KR or Flag-GLT-1 WT and immunoprecipitated with GLT-1 followed by immunoblotting for ubiquitin. Because previous work reported PMA induces GLT-1 ubiquitination (Sheldon et al., 2008), it was used to stimulate ubiquitination in this system. PMA treatment increased ubiquitinated GLT-1 in WT cells, whereas this effect was prevented in the cells expressing 7KR mutant (Supplemental Figure 6A).

**Figure 5.**
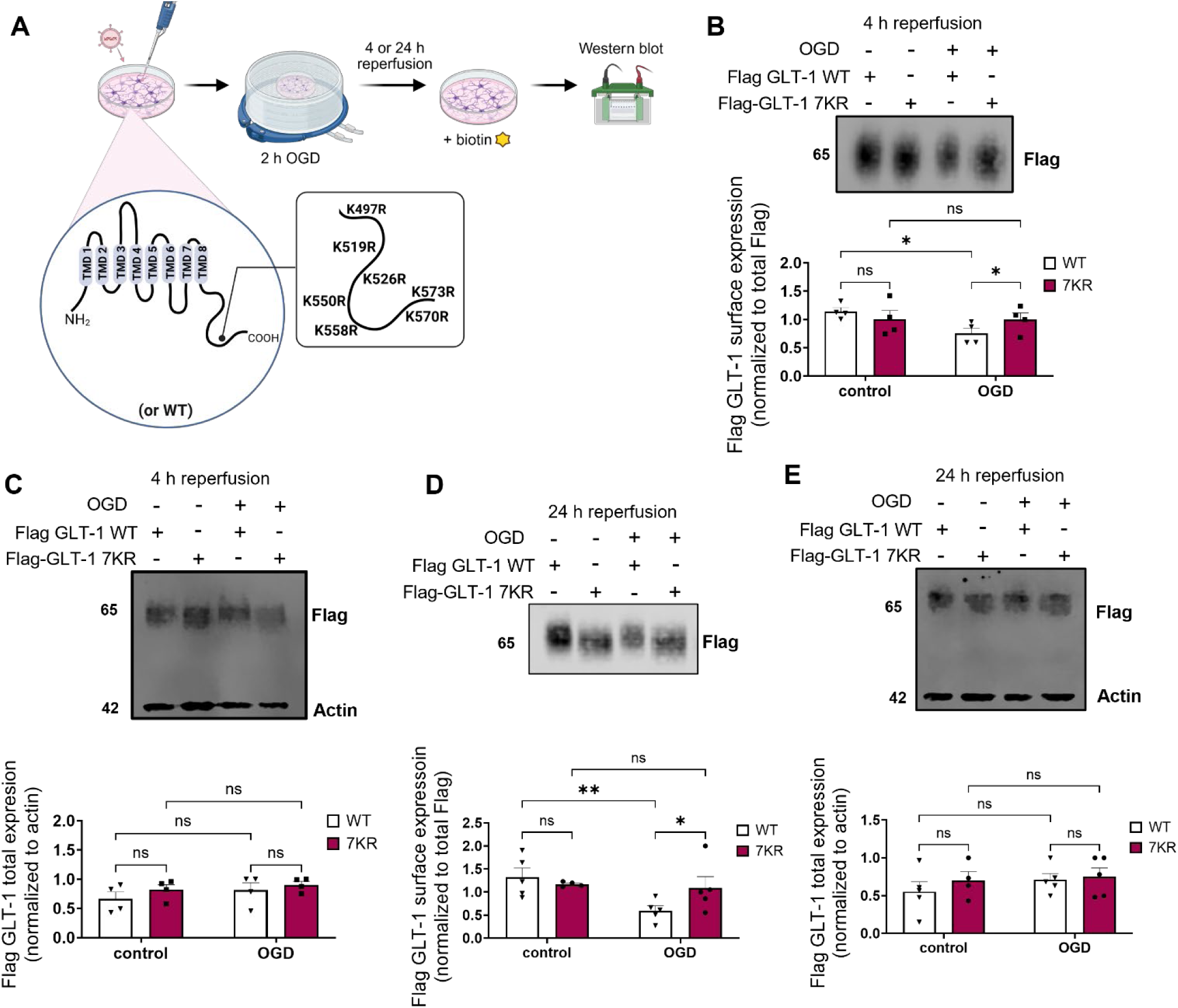
C-terminal lysine mutations stabilize GLT-1 surface expression after OGD. **A.** Schematic of experimental paradigm. Graphic displaying the C-terminal of GLT-1 (transmembrane domain 8, TMD8) depicting seven lysine (K) residues that were mutated to arginine (7KR). Glial cultures were transduced with lentivirus constructs encoding Flag-tagged GLT-1 WT or the C-terminal lysine mutant (Flag-GLT-1 7KR). Following OGD, cultures were biotinylated to measure transporter surface expression. **B-C**. Representative immunoblots (top), and protein quantification (bottom) of surface (**B**) and total (**C**) Flag-GLT-1 expression at 4 h reperfusion in WT- and 7KR- transduced cultures. **D-E**. Representative immunoblots (top), and protein quantification (bottom) of surface (**D**) and total (**E**) Flag GLT-1 expression at 24 h reperfusion in WT- and 7KR- transduced cultures. Data are presented as mean ± SEM, (n=4 independent culture preparations per group). Statistical analyses for data in panels B-C were performed using Two Way ANOVA followed by Fisher’s LSD *post-hoc* test. Data in panels D-E were performed using a mixed-effects model (REML) followed by Fisher’s LSD *post-hoc* test to account for missing value. * *p* < 0.05, ** *p* < 0.01, ns=non-significant.

Following validation, glial cultures were transduced with the Flag–GLT-1 WT or 7KR constructs 8 days prior to OGD, and surface biotinylation was performed at respective reperfusion time points. Efficient transduction was confirmed by visualization of EGFP fluorescence (Supplemental Figure 6B). Under control conditions, no differences in surface or total GLT-1 were observed in glia cultures transduced with these constructs and immunoblotted for Flag to measure construct GLT-1 expression. However, following OGD, the reduction in surface GLT-1 observed in Flag-GLT-1 WT cultures was prevented in cultures expressing Flag-GLT-1 7KR, at 4 h reperfusion (Figure 5B, *p* < 0.05 WT vs 7KR in OGD group) and 24 h reperfusion (Figure 5D, *p* < 0.05), with no changes in total GLT-1 expression (Figure 5C and 5E).

To further examine GLT-1 trafficking, GLT-1 colocalization with the early endosome marker EEA1 was analyzed by immunofluorescence via confocal imaging (Figure 6A). In Flag-GLT-1 WT-transduced cultures, OGD significantly increased GLT-1 colocalization with EEA1 (Figure 6B, *p* < 0.01 GLT-1 WT control vs GLT-1 WT OGD). This increase was prevented in cultures expressing Flag-GLT-1 7KR (Figure 6B, 7KR control vs 7KR in OGD group). These results suggest that PTMs at C-terminal lysine sites contribute to aberrant GLT-1 trafficking and internalization following OGD.

**Figure 6.**
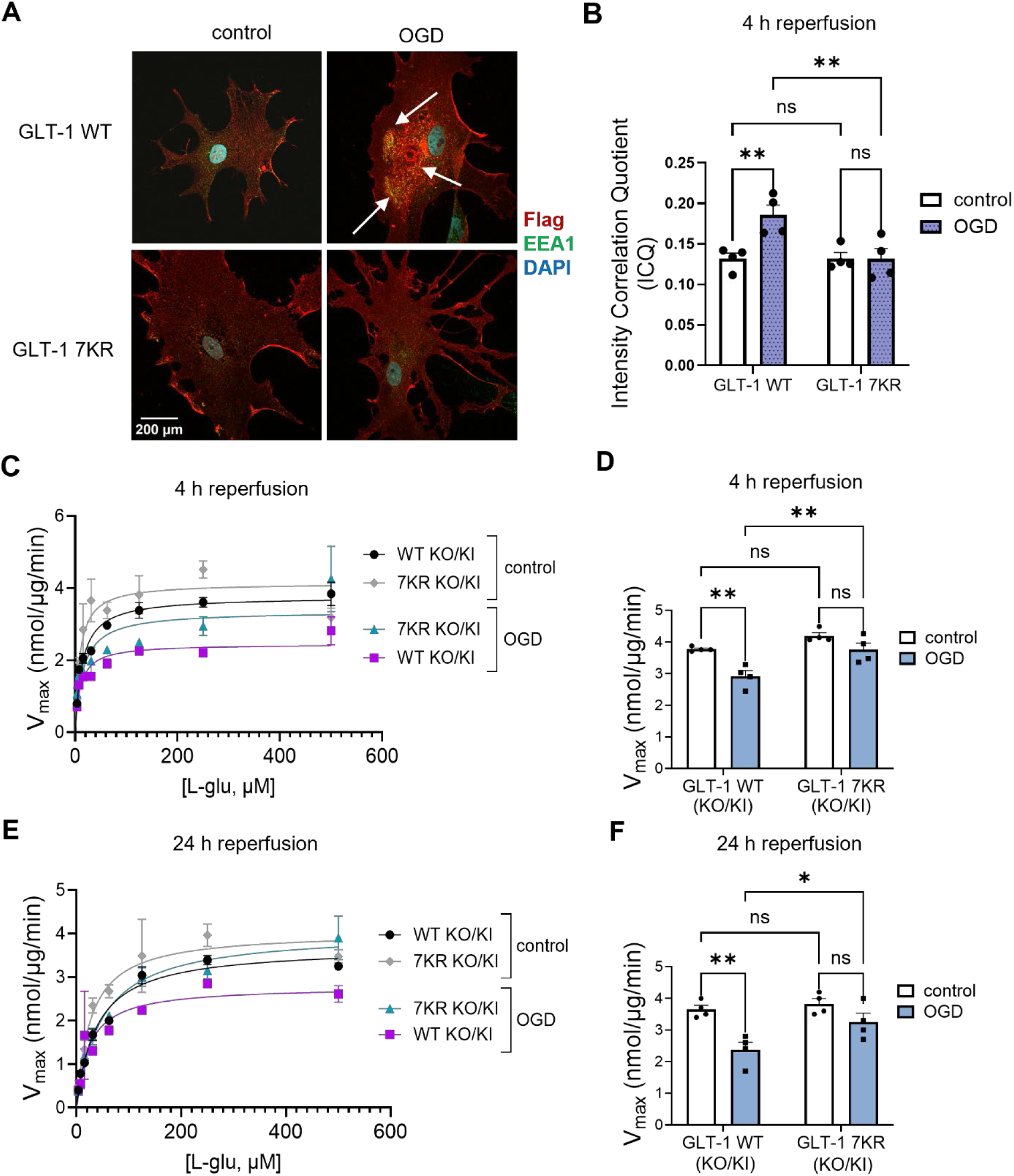
Inhibition of C-terminal lysine PTMs prevent GLT-1 internalization and restore functional uptake. **A.** Representative immunofluorescence images showing colocalization of Flag GLT-1 with the early endosome marker EEA1 following 4 h reperfusion. Arrows indicate regions of colocalization. Scale bar: 200 µm. **B**. Intensity Correlation Quotient (ICQ) analysis quantifying colocalization between Flag GLT-1 and EEA1 at 4 h reperfusion. **C-F**. Glutamate uptake assays were performed 4 h (**C, D**) or 24 h (**E, F**) after reperfusion in GLT-1-KO/KI 7KR or WT cultures**. C, E.** Representative Michaelis-Menten saturation curves of L-_3_H-glutamate uptake. **D, F.** Quantification of V_max_ (nmol/µg/min) for each condition (n=4 independent culture preparation per group with triplicates averaged). Data are presented as mean ± SEM. Statistical analyses were performed using two-way ANOVA followed by Fisher’s LSD *post-hoc* test. * *p* < 0.05, ** *p* < 0.01, ns= non-significant

Although these results demonstrated that C-terminal lysine modifications are drivers of GLT-1 downregulation following ischemic insult, it remained unclear whether their inhibition would translate into a functional recovery of transporter function. To assess uptake specifically from virally expressed GLT-1 constructs, we designed a lentiviral knockdown–knock in (KO/KI) system incorporating a miR30-based shRNA targeting endogenous GLT-1, allowing simultaneous knockdown of native GLT-1 and overexpression of either WT or 7KR GLT-1 (hereafter termed GLT-1-KO/KI WT and GLT-1 KO/KI 7KR, respectively). We first validated the GLT-1 miR30-based shRNA in COS-7 cells and demonstrated a 40-50% knockdown efficiency measured through glutamate uptake assays (Supplemental Figure 6C). Then, L-^3^H-glutamate uptake analysis was performed after OGD in glia cultures transduced with GLT-1-KO/KI WT or 7KR constructs at 4 h or 24 h reperfusion. Representative Michaelis-Menten curves are shown in Figure 6C and 6E. In GLT-1 KO/KI WT cultures, OGD significantly reduced V_max_ at both 4 h and 24 h reperfusion (Figure 6D and 6F, *p* < 0.01 WT control vs OGD). In contrast, this reduction was prevented in GLT-1 KO/KI 7KR cultures, with V_max_ significantly higher in than WT following OGD (Figure 6D and 6F, *p* < 0.01 at 4 h and *p* < 0.05 at 24 h, GLT-WT vs 7KR in OGD group). Notably, OGD did not significantly change V_max_ within the 7KR group (Figure 6D and 6F), and K_m_ values were unchanged across all conditions (4h: F (3,12) = 0.5862, *p* = 0.6355; 24 h: F(3,12) = 0.5719, *p* = 0.6442). Together, this data suggests that preventing the C-terminal lysine modifications stabilizes GLT-1 at the cellular surface and preserves glutamate uptake functionality following ischemic insult..

### 3.6 Ubiquitination Mediates Early GLT-1 Surface Downregulation following OGD

As mutation of the C-terminal lysine residues prevents multiple lysine-directed PTMs, including both ubiquitination and SUMOylation, we next asked whether specific inhibition of ubiquitination complexes is sufficient to drive the aberrant trafficking of GLT-1 following OGD. This approach was guided by our observation that GLT-1 ubiquitination, but not SUMOylation, increases at 4 h reperfusion, the time point when transporter internalization first occurs (Figure 1H). Additionally, pharmacological inhibitors of ubiquitination are more widely established than SUMOylation inhibitors, which we were unable to reliably validate in our system. To test this, cultures were immediately treated following OGD with TAK-243, an inhibitor of the E1 activation enzyme UBA1 (Ubiquitin-like modifier-activating enzyme 1, a broad E1 enzyme), which catalyzes the first step of the ubiquitination cascade (Lambert-Smith et al., 2020). Surface GLT-1 expression was then assessed by biotinylation and immunoblotting at 4 h reperfusion (Figure 7A). In basal/control conditions, TAK-243 treatment had no effect on surface or total protein levels of GLT-1; however, following OGD, TAK-243 treatment prevented the reduction in surface GLT-1 observed in vehicle-treated cultures (Figure 7B and 7C, *p* < 0.01 vehicle vs TAK in OGD group). Similar effects were observed at 24 h reperfusion (Supplemental Figure 7A and 7B), however it is important to note that TAK-243 application for this extended period of time resulting in a significant increase in total GLT-1 expression levels, supporting previous groups findings that transporter ubiquitination is mechanism of physiological turnover (Martínez-Villarreal et al., 2012).

**Figure 7.**
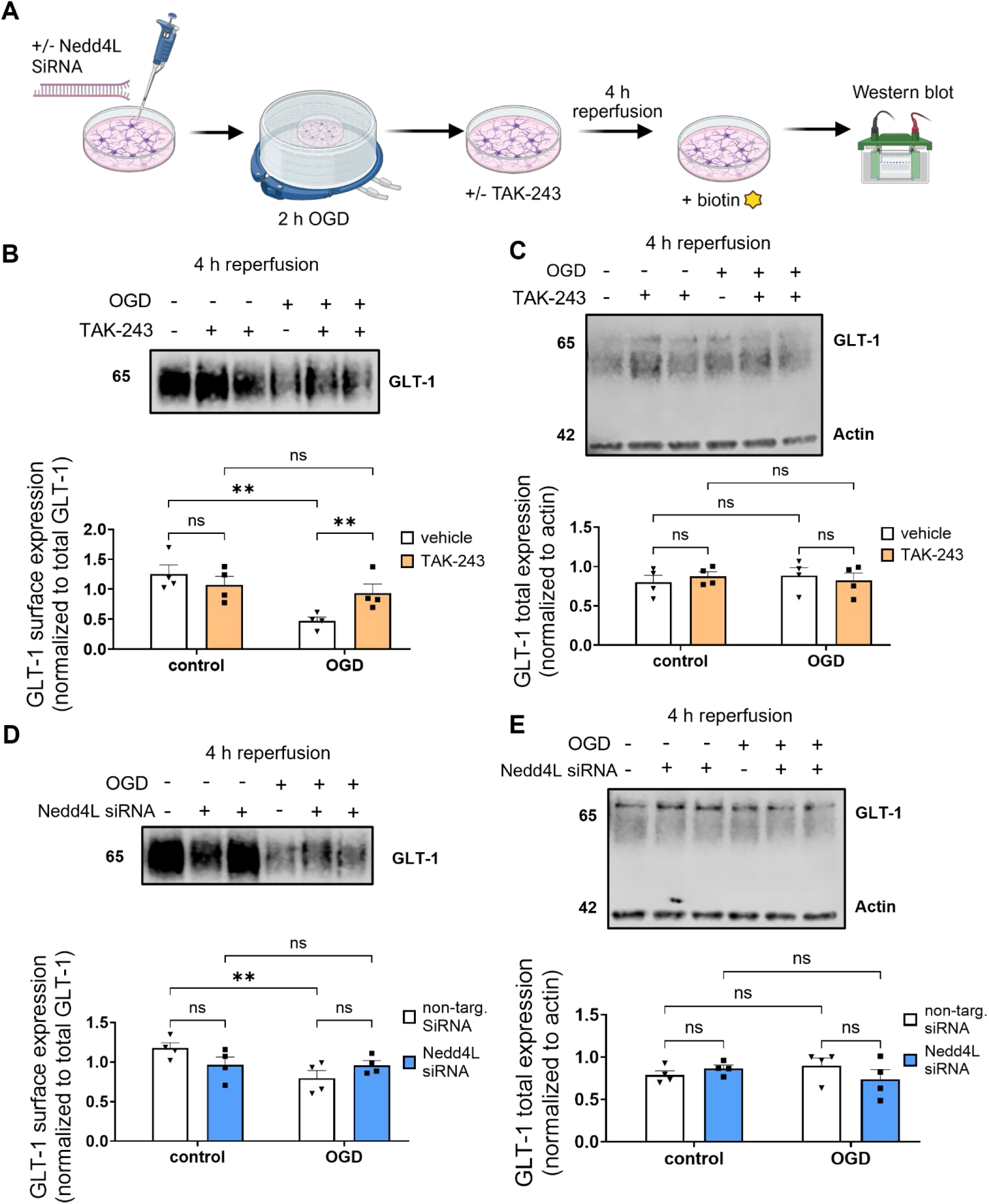
Inhibition of GLT-1 ubiquitination prevents surface downregulation following OGD. **A.** Schematic of experimental paradigm. A subset of cultures was transfected with Nedd4L siRNA and the other subset was treated with TAK-243 following OGD, followed by surface biotinylation and immunoblotting at the indicated timepoints. **B-C.** Representative immunoblots (top), and protein quantification (bottom) of surface (**B)** and total (**C**) GLT-1 expression following 4 h reperfusion with TAK-243 (100 nM) applied immediately after OGD. **D-E.** Glial cultures were transfected with Nedd4L siRNA 72 h prior to OGD. Representative immunoblots (top), and protein quantification (bottom) of surface (**D**) and (**E**) GLT-1 expression at 4 h reperfusion. Data are presented as mean ± SEM, (n=4 independent culture preparations per group). Statistical analyses were performed using Two Way ANOVA followed by Fisher’s LSD *post-hoc* test. ** *p* < 0.01, ns= non-significant.

Previous work has identified Nedd4L as an E3 ubiquitin ligase mediator of PKC-dependent ubiquitination and degradation of GLT-1 (Garcia-Tardon et al., 2012). Additionally, Nedd4L was also shown to mediate the downregulation of GLT-1 through activation of serum-dependent kinases (Boehmer et al., 2003). Based on these findings, we next investigated whether Nedd4L contributes to OGD-induced GLT-1 ubiquitination and downregulation following ischemic injury. Nedd4L was knocked down using targeted siRNA to assess its role in post-OGD GLT-1 trafficking. Validation of Nedd4L siRNA knockdown can be seen in Supplemental Figure 7C. While Nedd4L knockdown did not significantly increase GLT-1 surface expression relative to its vehicle counterpart following OGD, the reduction in surface GLT-1 observed between control and OGD conditions was no longer evident in Nedd4L siRNA-treated cultures (Figure 7D, Nedd4L siRNA control vs OGD, blue bars). Similar effects were observed at 24 h reperfusion (Supplemental Figure 7D and 7E). These findings suggest that Nedd4L contributes to OGD-induced downregulation of GLT-1 but is not solely responsible for its dysregulated trafficking.

### 3.7 Inhibition of GLT-1 C-terminal Lysine PTMs Confers Neuroprotection following OGD

To evaluate the neuroprotective potential of inhibiting ubiquitination-driven aberrant trafficking of GLT-1 following OGD, we employed an *ex vivo* organotypic slice culture (OSC) model to quantify cytotoxicity. Because our primary glial cultures consist predominantly of astrocytes with limited neuronal content (allowing for in depth analysis of astrocytic glutamate transporter regulation), OSCs were used to better model neuronal vulnerability to cellular death and preserve regional CNS architecture.

To confirm dysregulation of GLT-1 surface availability across model systems, we first quantified GLT-1 protein expression in both the cortex and hippocampus following a 30-minute OGD insult. In both regions, surface GLT-1 expression was significantly reduced following 4 h reperfusion (Figure 8B and 8C), and returned to baseline by 24 h reperfusion (Figure 8D and 8E). Based on the established vulnerability of the hippocampus to glutamate-excitotoxicity after ischemic insult (Butler et al., 2010; Caba et al., 2021), subsequent cytotoxicity experiments were performed in hippocampal OSCs.

**Figure 8.**
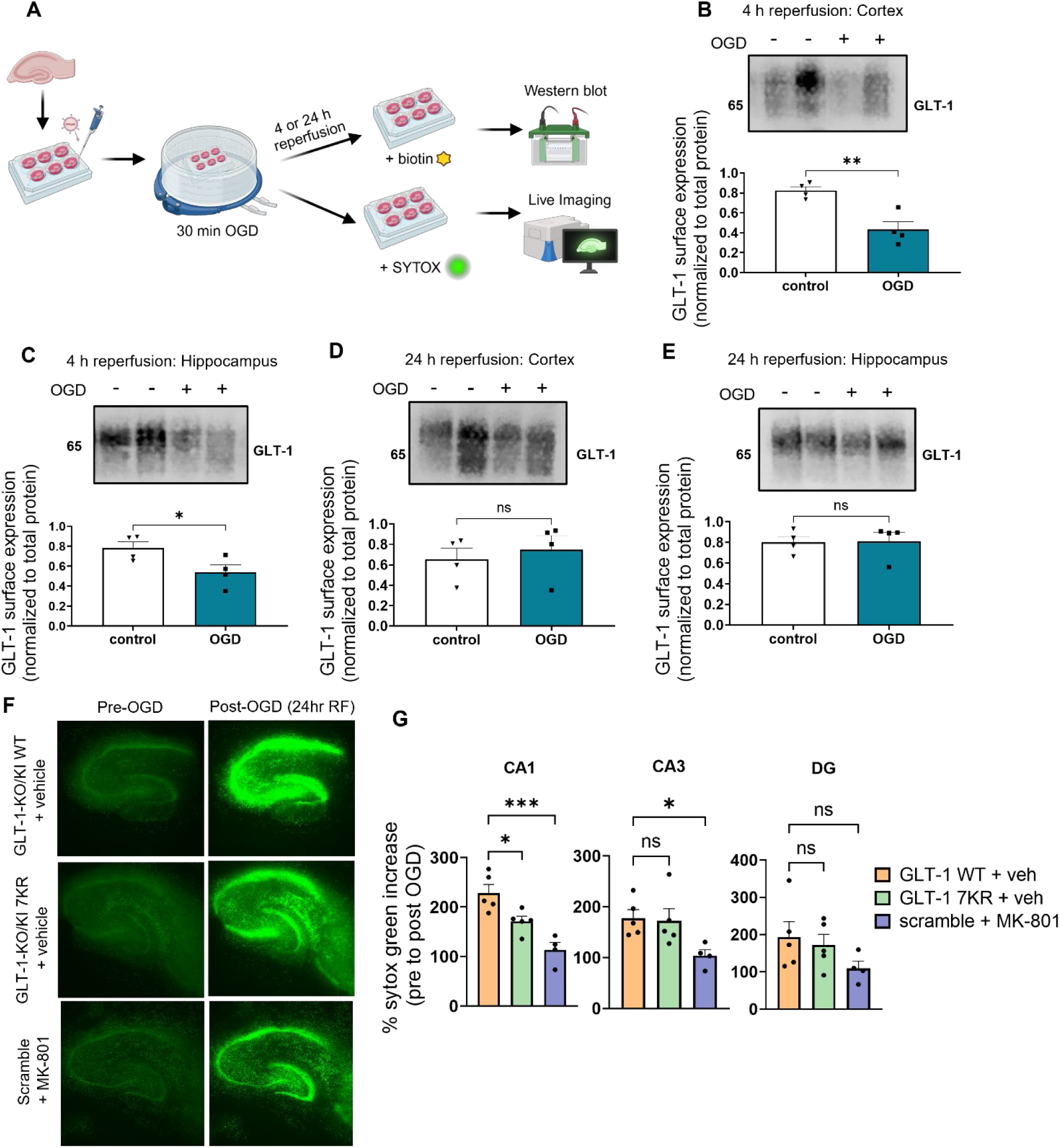
Inhibition of GLT-1 C-terminal lysine PTMs confers neuroprotection following OGD. **A.** Schematic of experimental paradigm. **B-C.** Representative immunoblot (top), and protein quantification (bottom) of surface GLT-1 expression in cortical (**B**) and hippocampal (**C**) organotypic slice cultures (OSCs) following 30 min OGD and 4 h reperfusion (n=4 independent OSC preparations per group; technical duplicates averaged). **D-E.** Representative immunoblots (top), and protein quantification (bottom) of surface GLT-1 expression in cortical (**D**) and hippocampal (**E**) OSCs following 30 min OGD and 24 h reperfusion (n=4 independent OSC preparations per group). **F.** Representative images of hippocampal OSCs transduced with GLT-1 KO/KI WT, GLT-1 KO/KI 7KR), or scramble virus and subjected to Sytox Green assay to assess cell death at 24 h reperfusion (10 days after transduction). The NMDA receptor antagonist MK-801 (10 µM) was applied during OGD and maintained throughout reperfusion as a positive control. **G.** Images were taken at a 4x objective. **F-G** Quantification of cell death in the CA1, CA3 and dentate gyrus following OGD (n=4-5 independent OSC preparations per group; technical triplicates averaged). Data are presented as mean ± SEM. Statistical analyses were performed using two-tailed unpaired *t* test or One Way ANOVA followed by Dunnett’s *post-hoc* test (compared to GLT-1 WT). * *p* < 0.05, ** *p* < 0.01, *** *p* < 0.001, ns=non-significant

To elucidate the neuroprotective potential of inhibiting GLT-1 ubiquitination in hippocampal OSCs, cell death was measured using SYTOX Green staining, a high-affinity green, fluorescent nucleic acid marker with penetration ability into comprised and damaged cells (Levraut et al., 2003; Truernit E, 2008). To avoid confounding neuroprotection from GLT-1 overexpression, which enhances extracellular glutamate clearance and limits excitotoxicity (Weller et al., 2008; Krzyzanowska et al., 2017), OSCs were transduced with the GLT-1 KO/KI WT or 7KR replacing endogenous GLT-1 with the respective viral constructs, or a scramble control. OSCs were transduced 24 h after slicing, and OGD was performed 9 days later. Adequate transduction was confirmed by mCherry fluorescence signal (Supplemental Figure 8A).

Background fluorescent images were acquired prior to OGD. Following OGD, slices were incubated with 100 µM SYTOX Green and imaged hourly for 24 h (Figure 8F). MK-801 (10 µM) was used as a post-treatment positive control. Compared to WT, slices expressing GLT-1 7KR exhibited significantly reduced cell death in the CA1 region following OGD, whereas no significant differences were observed in the CA3 and dentate gyrus (Figure 8G, *p* < 0.05 GLT-1 WT vs 7KR in CA1).

Collectively, these results indicate that preventing C-terminal lysine modifications of GLT-1 reduces neuronal injury following ischemic insult, supporting a role for GLT-1 ubiquitination as a mechanism driving aberrant transporter trafficking and excitotoxic damage. These results provide proof-of-concept that targeting lysine-directed post-translational modifications of GLT-1 may represent a neuroprotective strategy following ischemic injury.

## 4 Discussion

The downregulation of GLT-1 in response to ischemic injury has been widely reported (Raghavendra Rao et al., 2000; J.C. Chen et al., 2005; Krzyzanowska et al., 2014); however, the mechanisms that regulate its trafficking under ischemic conditions remain poorly defined. In this study, we identify a mechanism by which ischemic injury disrupts astrocytic glutamate transporter regulation. We demonstrate that OGD induces rapid internalization of GLT-1, leading to reduced transporter surface expression and impaired glutamate uptake. Mechanistically, this process is driven by lysine-directed PTMs of the GLT-1 C-terminal domain, particularly ubiquitination, which promotes transporter internalization and degradation. Preventing these modifications through mutating C-terminal lysine residues restores GLT-1 surface expression, preserves glutamate uptake, and reduces neuronal injury in organotypic slice cultures. Together, these findings identify ubiquitination-dependent GLT-1 trafficking as a key regulator of post-ischemic glutamate homeostasis.

First, we demonstrate that ischemic insult induces aberrant GLT-1 trafficking accompanied by reduced transport capacity. Following OGD, GLT-1 surface expression was downregulated, resulting in a decrease in V_max_. Notably, changes in surface GLT-1 protein levels were not observed until 4 h reperfusion but remained reduced for up to 24 h. This temporal profile is consistent with previous literature suggesting that GLT-1 has a half-life exceeding 24 h (Sheldon et al., 2008; Martinez-Lozada et al., 2016).

Reduced surface GLT-1 could be a result of two primary mechanisms: an increase in the internalization of GLT-1, or a decreased insertion of newly synthesized protein. Inhibition of dynamin with Dynasore prevented downregulation of surface GLT-1, therefore pointing to the internalization of the transporter as the primary mechanism in response to OGD. Following the internalization of GLT-1 there was an increase in degradation. Interestingly, inhibiting either the lysosomal or the proteasomal pathway alone was insufficient to prevent GLT-1 downregulation, whereas combined inhibition of both pathways preserves transporter expression. This could either suggest that GLT-1 degradation in response to ischemic injury occurs through multiple pathways, or that inhibition of one degradation route triggers compensatory degradation through the other. Although ubiquitination was initially characterized as a signal for proteasomal degradation, emerging evidence suggests that ubiquitin-tagged proteins can be routed through proteasomal processing pathways while ultimately being degraded in the lysosome (Alwan et al., 2003). This phenomenon may therefore explain why combined treatment with Bafilomycin and MG-132 most effectively preserved GLT-1 expression levels. Together, these data demonstrate that GLT-1 undergoes degradation following ischemic injury, although the precise trafficking pathway needs to be further studied.

We next sought to determine molecular drivers of GLT-1 internalization and degradation following OGD. Previous work has demonstrated the important role that post-translational modifiers play in the regulation of GLT-1 trafficking. The constitutive turnover of GLT-1 was found to be dependent on ubiquitination/deubiquitination cycles that ensure appropriate transporter availability (Martínez-Villarreal et al., 2012). Notably, the ubiquitination and subsequent degradation of GLT-1 has been found to be a major driver of GLT-1 downregulation in a model of Parkinson’s disease (Zhang et al., 2017) and the SUMOylation of GLT-1 as a driver of transporter compartmentalization (Foran et al., 2014). We therefore sought to identify if any of these modifiers also played a role in GLT-1 trafficking following OGD. In the present study, both GLT-1 ubiquitination and SUMOylation were increased following OGD, but with different temporal profiles. We found an increase in GLT-1 ubiquitination at the 4 h reperfusion timepoint, coinciding with the onset of GLT-1 internalization, and returned to baseline levels by 24 h reperfusion, whereas SUMOylation increased at later time points. This temporal separation suggests that distinct lysine-directed PTMs may regulate different stages of GLT-1 trafficking during ischemic insult, with ubiquitination mediating early transporter internalization and SUMOylation potentially contributing to later regulatory processes. Pharmacological ubiquitination inhibitors prevented GLT-1 downregulation, suggesting ubiquitination as a main driver of early post-ischemic GLT-1 trafficking. GLT-1 internalization has previously been shown to occur in an activity-dependent manner in response to elevated extracellular glutamate, which can promote recruitment of arrestin and E3 ubiquitin ligases of the Nedd4 family to the transporter, resulting in ubiquitination and subsequent endocytosis of GLT-1 from the plasma membrane surface (Garcia-Tardon et al., 2012; Ibáñez et al., 2016). While this mechanism likely serves to regulate transporter availability under physiological conditions, ischemia-induced glutamate accumulation may pathologically amplify this process and contribute to sustained loss of surface GLT-1. However, these findings do not exclude involvement of GLT-1 SUMOylation in later phases of transporter regulation. Interestingly, a previous study found that the SUMOylation of proteins can antagonize ubiquitin-dependent proteasome pathway by competing for shared lysine residues; for example, the SUMOylation of IκB prevents its ubiquitination and subsequent proteasomal degradation and inhibition of NF-κB transcription (Desterro et al., 1998). A similar competitive mechanism could explain the bidirectional alterations between SUMOylated and ubiquitinated GLT-1 observed here and may suggest that the SUMOylation is a protective mechanism in limiting further GLT-1 degradation following ischemic injury.

The C-terminal domain of GLT-1 has been shown to be a primary domain of post-translational modifications that mediate transporter trafficking. Specifically, the ubiquitination of the 7 lysine residues of the transporters C-terminal domain was shown to be sufficient to drive the internalization of the GLT-1 through a PKC-mediated manner (Gonzalez-Gonzalez et al., 2008; Sheldon et al., 2008). Importantly, these C-terminal lysine residues are also important sites for transporter SUMOylation. Although SUMOylation commonly occurs within the canonical sequence ψKxE, where ψ is an aromatic amino acid, of which the C-terminal domain of GLT-1 has a single site (Da Silva-Ferrada et al., 2012), prior *in vitro* work found that SUMOylation of GLT-1 was only abrogated completely when all lysine residues of the C-terminal tail were unable to be modified (Foran et al., 2014), suggesting noncanonical modification sites. Therefore, it is also important to consider that SUMOylation or ubiquitination may occur at shared lysine residues. With this, we designed a lentiviral construct that would allow us to study the role of lysine-mediated post-translational modifications in the trafficking of GLT-1 following ischemic insult. Our results show that by preventing PTMs interactions of GLT-1 C-terminal lysine residues, GLT-1 surface availability is restored after OGD at both 4 and 24 h reperfusion, which suggests that lysine-directed PTM interactions drive transporter downregulation. Furthermore, blockade of these modifications rescued uptake velocity, due to bolstering of GLT-1 at the cell surface. Together, these findings suggest a model in which aberrant ubiquitination and/or SUMOylation of GLT-1 following OGD reduces transporter availability and impairs glutamate clearance.

To further probe whether SUMOylation or ubiquitination primarily mediates ischemia-induced alterations in GLT-1 trafficking, we used a pharmacological approach to selectively inhibit these interactions. We focused primarily on ubiquitination, as OGD-induced GLT-1 ubiquitination temporally coincides with transporter internalization. Inhibition of UBA1 a central protein in the ubiquitination cascade, preserved surface GLT-1 expression following OGD, supporting ubiquitination as a principal driver of early ischemia-induced GLT-1 downregulation. Although C-terminal lysine mutation prevents both ubiquitination and SUMOylation, the delayed onset of SUMOylation relative to ubiquitination and internalization suggests that SUMOylation is likely to contribute to later regulatory events rather than initiating acute transporter loss. One limitation of the study is that pharmacological inhibition of UBA1 blocks global ubiquitination and therefore may affect additional cellular processes beyond GLT-1 regulation. However, the concordance between pharmacological inhibition and the GLT-1 7KR mutant supports a direct role for ubiquitination in regulating GLT-1 trafficking.

Further, Nedd4L has been identified as a key mediator of PKC-dependent GLT-1 ubiquitination and trafficking, and increased Nedd4L-mediated ubiquitination promotes GLT-1 degradation in a model of Parkinson’s disease (Zhang et al., 2017). Given the important role Nedd4L plays in GLT-1 trafficking, we wanted to determine its role in post-ischemic internalization of GLT-1. Our results show Nedd4L knockdown attenuated OGD-induced GLT-1 downregulation, although it did not significantly change surface transporter expression levels relative to vehicle-treated OGD cultures. This suggests that Nedd4L plays a mediatory but not exclusive role in the downregulation of GLT-1 following ischemic injury. Indeed, GLT-1 lacks canonical binding motifs for Nedd4 family ligases, and previous work has suggested that adaptor proteins such as caveolin-1 may facilitate recruitment of ubiquitin ligases to the transporter (Garcia-Tardon et al., 2012). Thus, even though Nedd4L contributes to GLT-1 downregulation following OGD, our findings may suggest that additional E3 ligases may participate in regulating GLT-1 ubiquitination under ischemic conditions and future work should focus on identifying these E3 ligase(s).

During preparation of this manuscript, Zhang et al. (2025) reported that GLT-1 ubiquitination contributes to transporter degradation following ischemic injury through regulation of the E3 ligase SMURF1. Consistent with these findings, our data also identify ubiquitination as an important regulator of GLT-1 trafficking following ischemic insult. However, the present study extends these observations by demonstrating that lysine-directed PTMs of the GLT-1 C-terminal domain directly control transporter internalization and degradation, providing a more comprehensive mechanistic understanding into how these PTMs regulate GLT-1 trafficking during ischemic stress.

As our C-terminal mutant was able to prevent OGD-induced downregulation of GLT-1 and restore functional uptake, we aimed to determine if this approach could also provide neuroprotection in response to ischemic challenge. To address this idea appropriately, we utilized hippocampal OSCs as this model system would allow us to first more accurately assess excitotoxic cell death in a tissue-intact system and secondly would provide a complete cell population profile as our primary glia cultures do not contain enough neurons to effectively measure cellular death.

Hippocampal OSCs transduced with GLT-1 KI/KO WT or 7KR constructs were subjected to OGD and stained with Sytox Green to assess cell death. Treatment with the NMDA antagonist MK-801 was able to reduce cell death in the CA1, CA3, and DG of the hippocampus. Promisingly, we observed reduced cell death in GLT-1 KI/KO 7KR transduced cultures compared to their WT counterparts, however, this was only observed in the CA1. Interestingly, previous work has shown that the CA1 is a primary region of hippocampal GLT-1 downregulation after ischemic injury (Zhang et al., 2019). Further, in a GLT-1 knockdown mouse line, the CA1 was found to be more susceptible glutamate neurotoxicity compared to other regions of the hippocampus (Tanaka et al., 1997). This could explain why our 7KR mutant was effective in preventing OGD-induced cell death only in the CA1 region of the hippocampus. These findings suggest that GLT-1 trafficking mechanisms may contribute to region-specific patterns of neuronal vulnerability during ischemia. A few limitations of our study should be noted, such as that the knockdown–knock-in system used in this study achieved only partial suppression of endogenous GLT-1 expression. While this approach allowed selective analysis of virally expressed constructs, residual endogenous transporter expression may still contribute to measured uptake. Although our findings demonstrate a clear role for lysine-directed PTMs in regulating GLT-1 trafficking following ischemic insult, these experiments were performed *in vitro* and in *ex vivo* cultures. The OGD model reproduces key metabolic features of ischemia but does not fully recapitulate the complex cellular and vascular interactions present during stroke *in vivo*. Thus, future studies using *in vivo* models of ischemic stroke will be required to determine whether the same mechanisms regulating GLT-1 trafficking are recapitulated in the intact brain.

In summary, our results demonstrate that ischemia induces aberrant trafficking of GLT-1, which results in impaired glutamate uptake. Because extracellular glutamate can remain elevated for up to 24 h following ischemic injury (Davalos et al., 1997), reduced transporter function may contribute to excitotoxic damage and hinder post-ischemic cellular repair. By targeting C-terminal lysine PTMs of GLT-1, we were able to prevent GLT-1 internalization and thereby increase functional uptake. Pharmacological inhibition of ubiquitination, a dominant lysine-mediated PTM that regulates GLT-1 trafficking, similarly prevented OGD-induced downregulation of the transporter, supporting ubiquitination as a principal driver of early ischemia-induced GLT-1 downregulation. However, since the C-terminal lysine mutations also block other lysine-directed modifications such as SUMOylation, we cannot exclude contributions from additional PTMs. Nonetheless, targeting C-terminal lysine post-translational modifications of GLT-1 conferred neuroprotection in hippocampal OSCs. While future studies are required to determine whether modulation of these pathways can be leveraged therapeutically *in vivo*, these findings provide proof of concept that preventing ischemia-induced aberrant GLT-1 trafficking could prove to be a promising therapeutic strategy.

## Author Contributions

**S. K. Gill:** Writing- original draft, conceptualization, study design, methodology, validation, data collection, formal analysis, visualization. **K. L. Reeb**: Methodology, reviewing and editing manuscript. **Kroll**: Data collection, reviewing and editing manuscript. **O. V. Mortensen**: Resources, study design, methodology, reviewing and editing manuscript. **A. C. K. Fontana**: Funding acquisition, resources, supervision, study design, reviewing and editing manuscript.

## Acknowledgments

We would like to thank Dr. Susan Amara (NIH) for providing the EAAT2 antibody and Dr. Michael Robinson (University of Pennsylvania) for the GLT-1 cDNA. I would also like to thank Dr. Antonio Sanz-Clemente, Dr. Joshua Jackson, Dr. Liang Qiang, and Dr. Elena Blanco-Suarez for their helpful discussion.

## Data Availability Statement

The data that supports the findings of this study are available in Supporting Information of this article.

## Abbreviations

aCSF: artificial cerebrospinal fluid
EEA1: Early Endosome Antigen 1
EAAT: excitatory amino acid transporter 1
EAAT2: excitatory amino acid transporter 2
CNS: central Nervous System
DIV: days in vitro
DMEM: Dulbecco’s Modified Eagle Medium
D-PBS: Dulbecco’s phosphate-buffered saline
GLT-1: Glutamate transporter-1
GLAST: Glutamate-aspartate transporter
GFAP: Glial fibrillary acidic protein
ICC: Immunocytochemistry
LDH: lactate dehydrogenase
MOI: Multiplicity of Infection
OSC: Organotypic slice culture
PBS-T: Dulbecco’s-Phosphate buffer with the addition of 0.05% Tween-20
PBS-CM: Dulbecco’s Phosphate buffer with addition of 0.1 mM CaCl_2_ and 1 mM MgCl_2_
PFA: Paraformaldehyde
PTM: Post-translational modification
RIPA: Radioimmunoprecipitation assay buffer
SEM: standard error of the mean
TNE: Tris-NaCl-EDTA

