## Supplemental File for "Ischemia-Induced Post-Translational Modifications of GLT-1 Mediate Aberrant Trafficking and Impaired Glutamate Uptake"

**Corresponding author:**

*Andréia C. K. Fontana

ORCID: 0000-0002-4791-8746

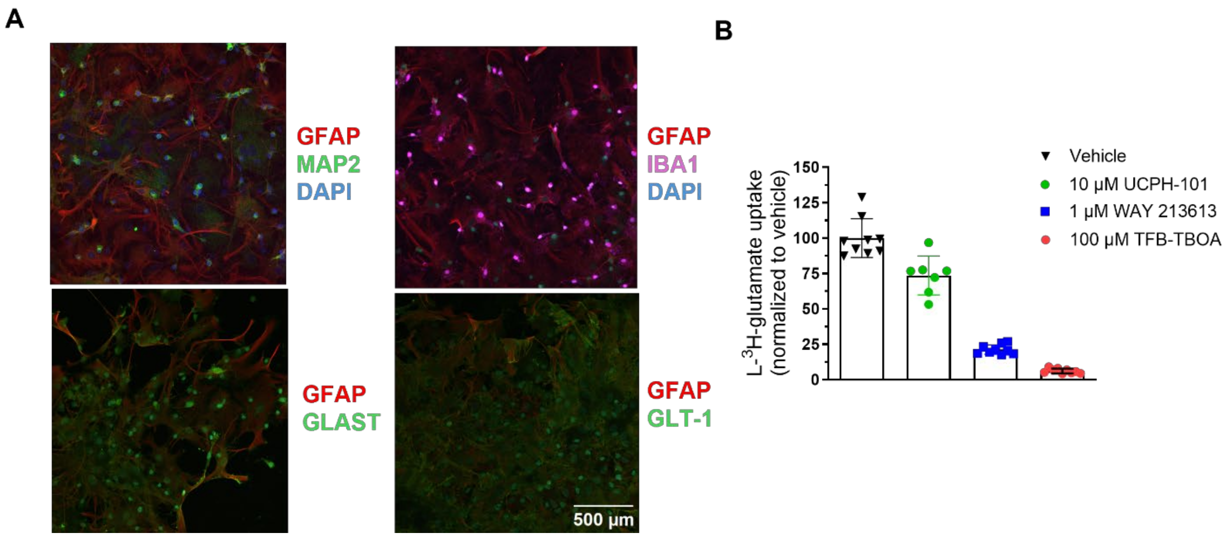


**Supplemental Figure 1. Cellular composition and glutamate transporter expression in primary glial cultures.**

**A**. Representative confocal immunofluorescence images showing cellular composition and transport expression in glia cultures. Top left: astrocyte marker GFAP and neuronal marker MAP-2. Top right: astrocyte marker GFAP and microglial marker IBA1. Bottom left: astrocyte marker GFAP and GLAST. Bottom right: GFAP and GLT-1. Scale bar: 500 µm. **B.** Quantification of glutamate uptake in presence of 1 µM of the selective GLAST inhibitor UCPH-101, the selective GLAST/GLT-1 inhibitor TFB-TBOA, and the selective GLT-1 inhibitor WAY 213613. Data are expressed as % of L-$^{3}$H-glutamate normalized to vehicle.

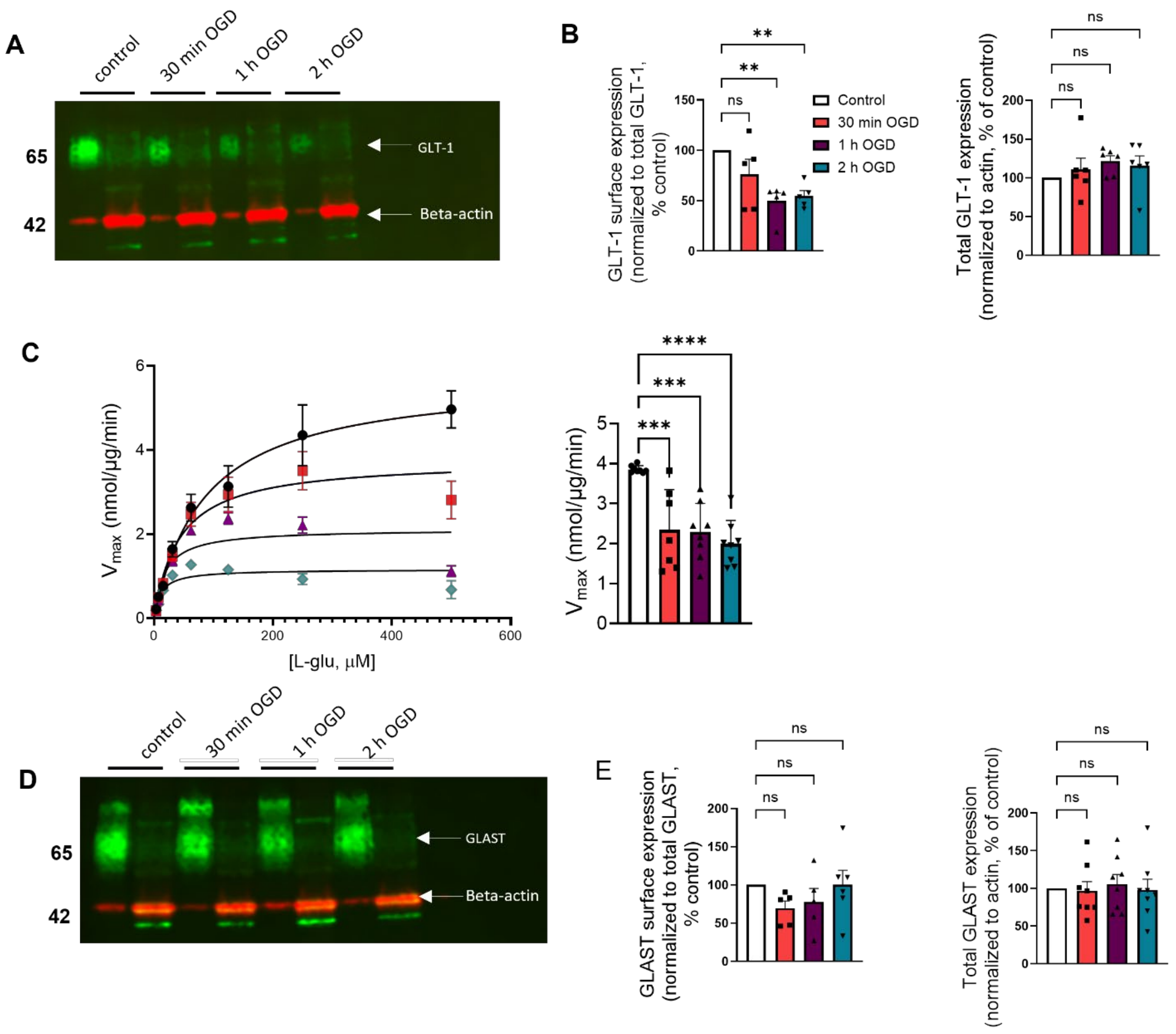


**Supplemental Figure 2. Increasing OGD duration reduces GLT-1 surface expression and transport velocity.**

**A.** Representative immunoblots showing GLT-1 surface and total expression (~65 kDa) following 30 minutes, 1 h, or 2 h OGD. **B.** Quantification of surface (left) and total (right) GLT-1 expression across varying OGD insult lengths (n=5-6 independent cell culture preparations per group). **C.** Representative Michaelis-Menten saturation curves of L-$^{3}$H-glutamate uptake measured at varying OGD lengths (right) and quantification of $V_{max}$ values normalized to nmol/µg/min (n=7 independent cell culture preparations per group; 11 replicates averaged per experiment). **D**. Representative immunoblots showing GLAST surface and total expression (~65 kDa) following 30 minutes, 1 h, or 2 h OGD. **D.** Quantification of surface (left) and total (right) GLAST expression across varying OGD insult lengths (n=5-6 independent cell culture preparations per group). Data are presented as mean ± SEM. Statistical analyses were performed using One Way ANOVA followed by Dunnett's multiple comparison *post-hoc* test. ** $p < 0.01$, *** $p < 0.001$, **** $p < 0.00001$, ns= non-significant.

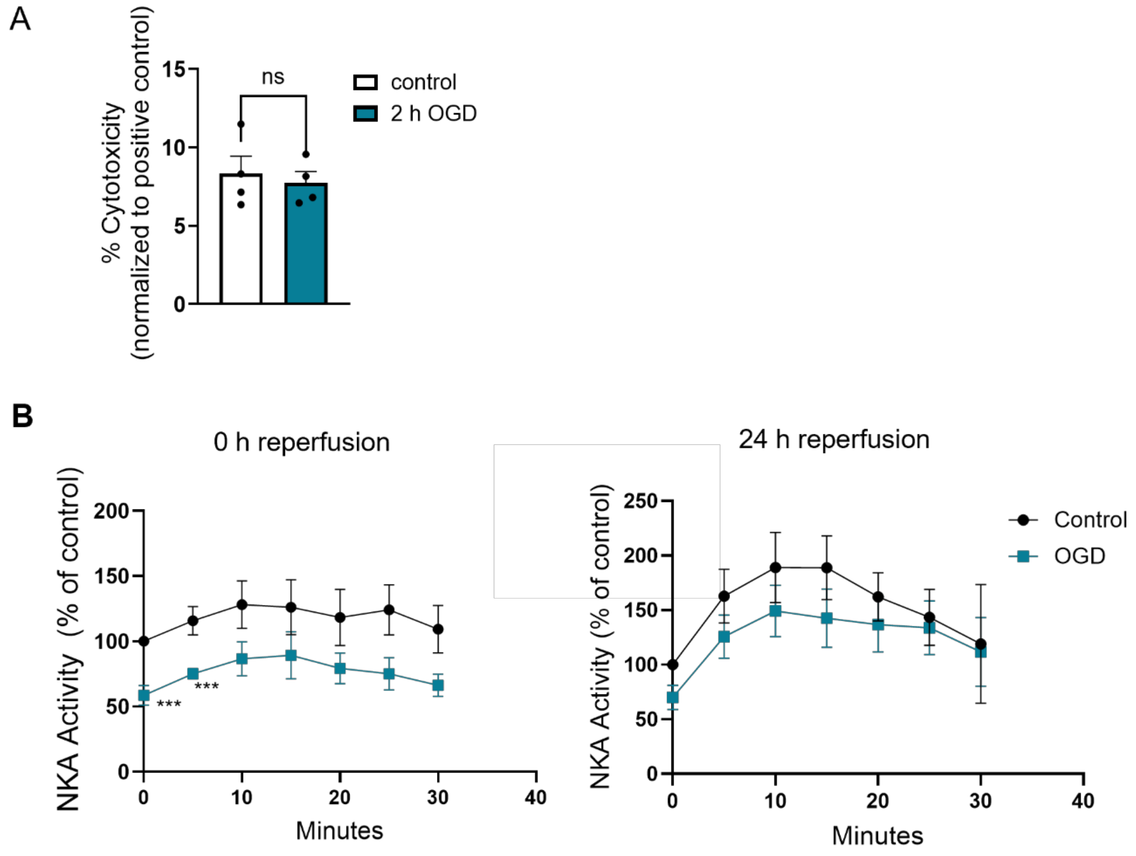


**Supplemental Figure 3. OGD does not induce cytotoxicity in glia cultures and transiently reduces $Na^+/K^+$-ATPase activity.**

**A.** Lactate dehydrogenase assay measuring level of cytotoxicity in glia cultures following OGD (n= 4 from independent cell cultures per group with six replicates averaged) **B.** Enzymatic activity of the NKA measured through a phosphatase fluorometric assay over a period of 35 min immediately following OGD (left) or after 24-hour reperfusion (right) (n= 5 from independent cell cultures per group with duplicates averaged). Data is presented as mean ± SEM. Statistical analyses were performed using One Way ANOVA followed by Dunnett's multiple comparison *post-hoc* test. *** $p$ < 0.001, ns= non-significant.

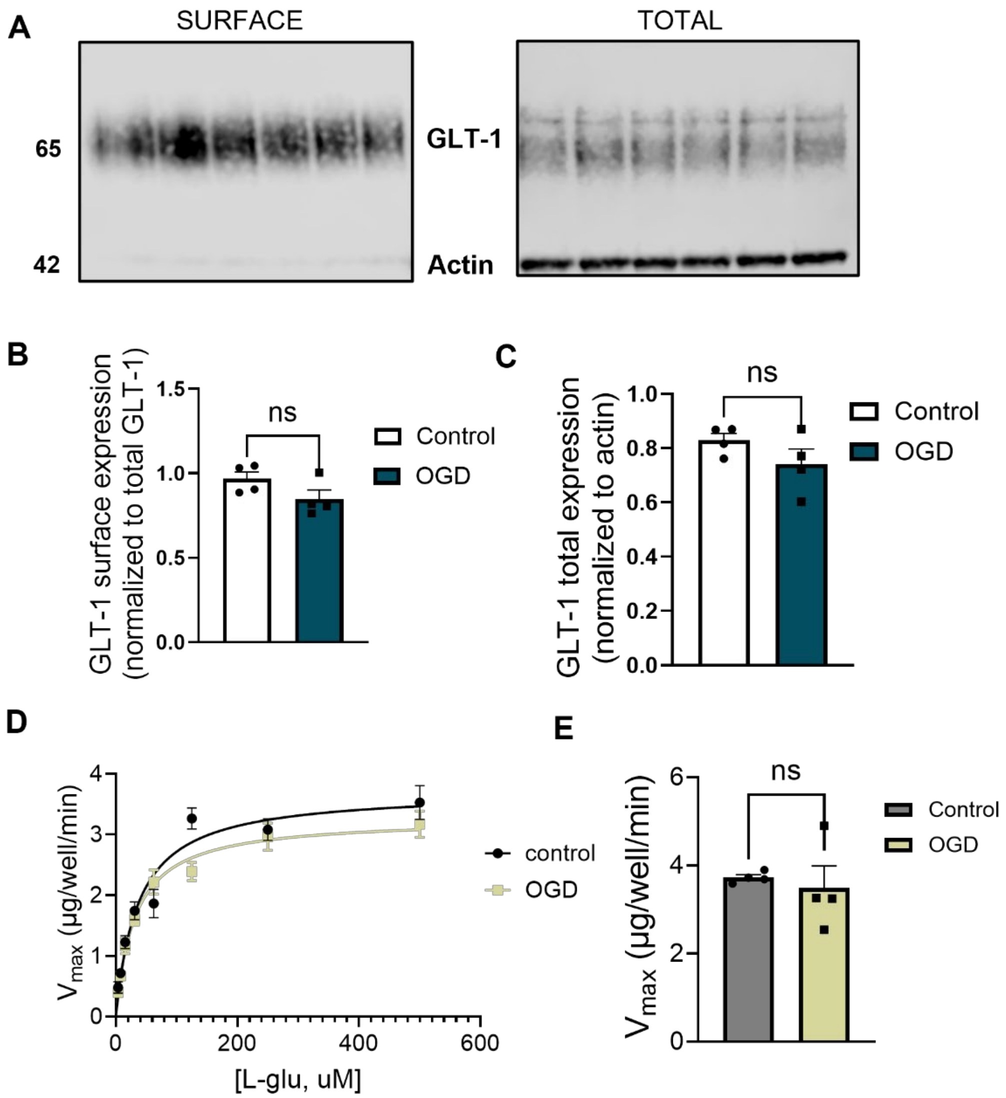


**Supplemental Figure 4. GLT-1 surface expression and glutamate transport velocity recover by 48 h reperfusion following OGD.**

**A.** Representative immunoblots showing GLT-1 surface and total expression (~65 kDa) following 2 h OGD and 48 h reperfusion. **B-C.** Quantification of surface (**B**) and total (**C**) GLT-1 expression following 2 h OGD and 48 h reperfusion (n=4 independent cell culture preparations per group; triplicates wells averaged). **D**. Representative Michaelis-Menten saturation curves of L-$^{3}$H-glutamate uptake measured at 48 h reperfusion. **E**. Quantification of $V_{max}$ values normalized to nmol/µg/min following 48 reperfusion (n=4 independent cell culture preparations per group; 11 technical replicates averaged per experiment). Data are presented as mean ± SEM. Statistical analyses were performed using two-tailed unpaired t tests. ns= non-significant.

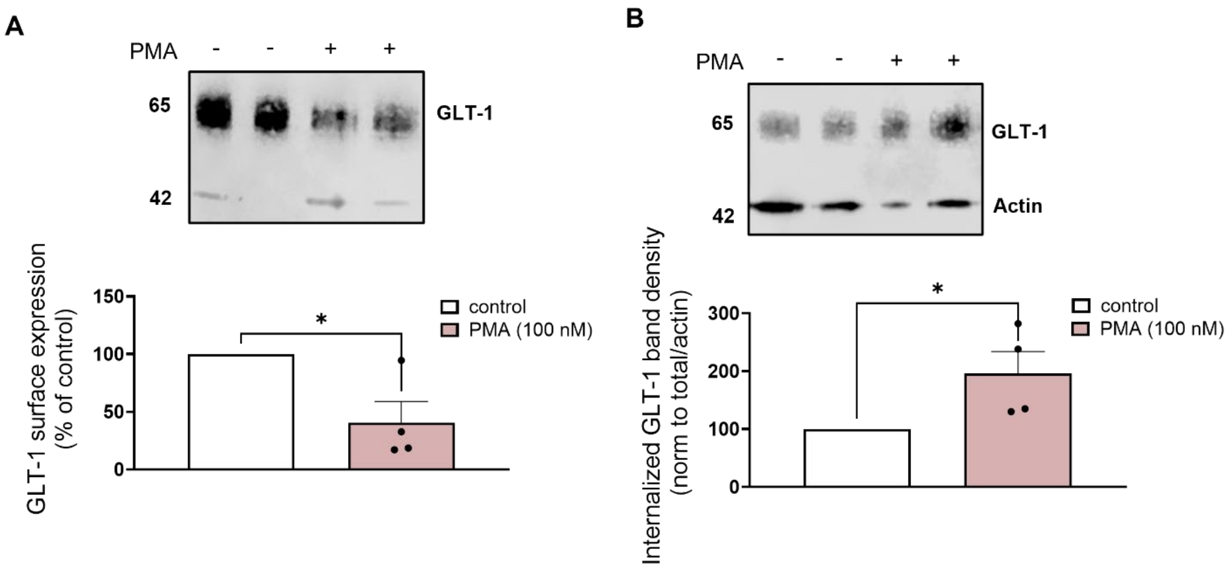


**Supplemental Figure 5. Phorbol 12-myristate 13-acetate (PMA) induces GLT-1 internalization.**
**A.** Representative immunoblots (top) and protein quantification (bottom) or internalized, pre-labeled GLT-1 measured by endocytic biotinylation following treatment with 100nM PMA. **B**. Representative immunoblot (top) and protein quantification (bottom) of total pre-labeled GLT-1 (remaining surface and internalized pools) following treatment with 100 nM PMA (n=4 independent cell culture preparations per group; technical duplicates averaged). Data are presented as mean ± SEM. Statistical analyses were performed using a two-tailed unpaired t test. * $p < 0.05$.

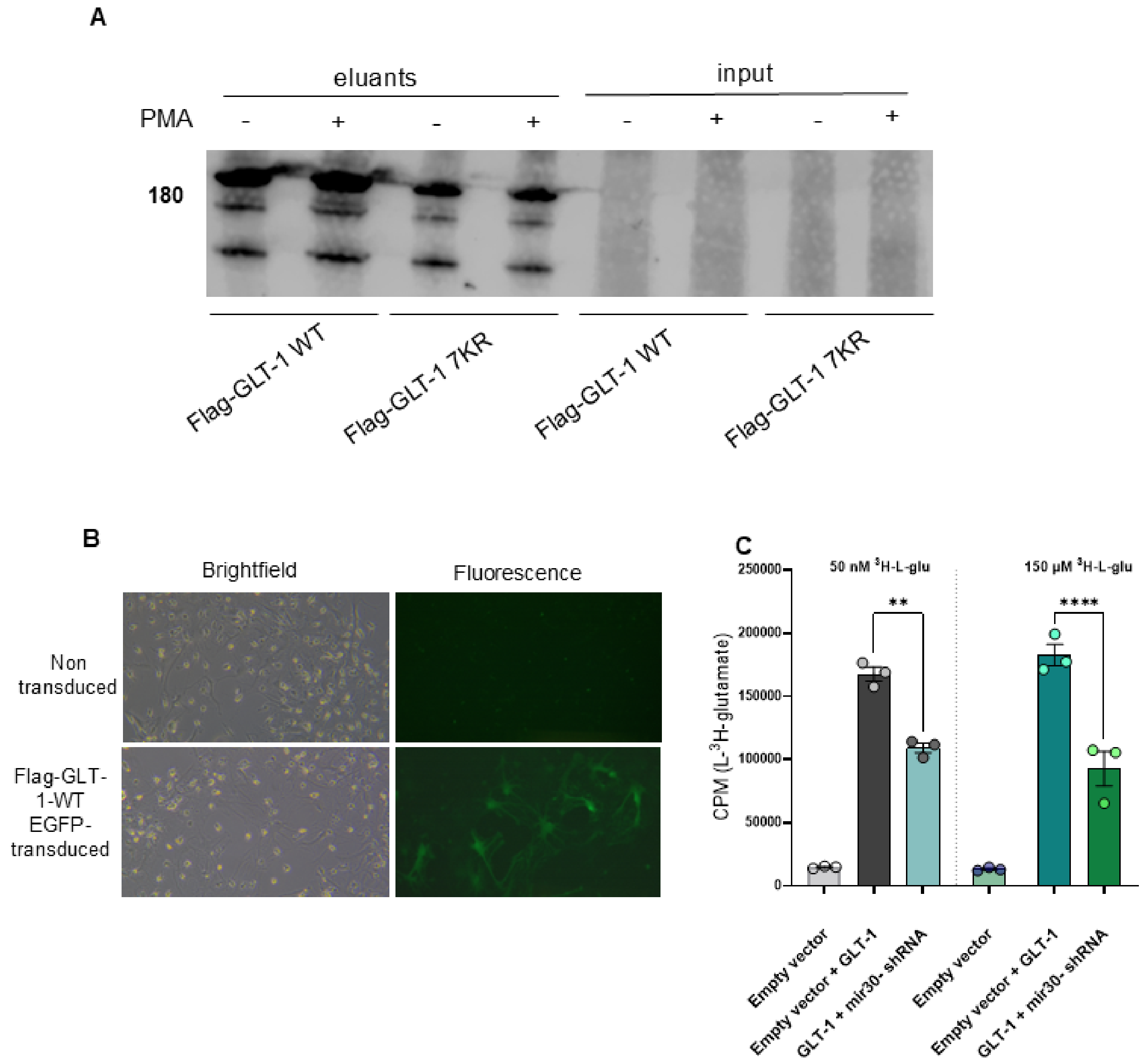


**Supplemental Figure 6. Validation of GLT-1 viral constructs and miR30 knockdown.**
**A.** Flag-tagged GLT-1 (WT or 7KR) was immunoprecipitated from COS-7 cell lysates and immunoblotted for ubiquitin (P4D1) to assess whether the 7KR construct effectively impeded GLT-1 ubiquitination **B**. Representative immunofluorescence images of glia cultures transduced with Flag-GLT-1 WT, compared to non-transduced cells, visualized using a GLF filter cube to detect EGFP fluorescence. **C**. Quantification L-$^{3}$H-glutamate uptake measured in COS-7 cells after transfection with empty vector, empty vector + GLT-1, or GLT-1 + mir30 siRNA, demonstrating knockdown efficiency of endogenous GLT01. Data are presented as mean ± SEM. Statistical analyses were performed using a One Way ANOVA followed by Sidaks multiple comparison *post-hoc* test. ** $p < 0.01$, **** $p < 0.0001$.

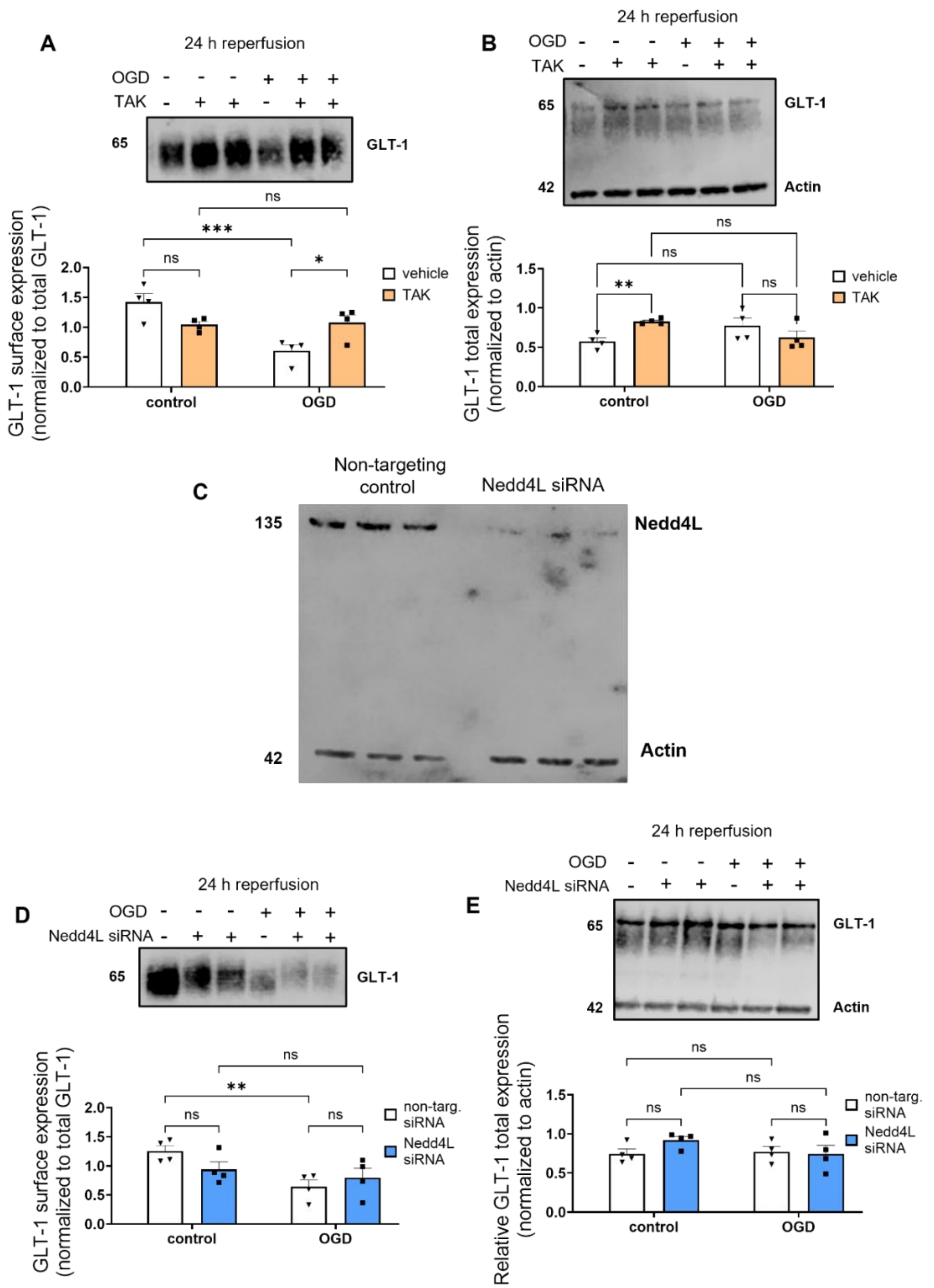


**Supplemental Figure 7. Inhibition of GLT-1 Ubiquitination Prevents OGD-induced Surface Downregulation.**

**A-B**. Representative immunoblots (top), and protein quantification (bottom) of surface (**A**) and total (**B**) GLT-1 expression following 24 h reperfusion with TAK-243 (100 nM) applied immediately after OGD. **C.** Representative immunoblot showing Nedd4L expression 72 h after transfection with Nedd4L siRNA or non-targeting control. **D-E**. Glial cultures were transfected with Nedd4L siRNA 72 h prior to OGD. Representative immunoblots (top), and protein quantification (bottom) of surface (**D**) and (**E**) GLT-1 expression at 24 h reperfusion. Data (**A-B, D-E**) are presented as mean ± SEM (n=4 independent culture preparations per group). Statistical analyses were performed using Two Way ANOVA followed by Fisher's LSD *post-hoc* test. ** $p < 0.01$, *** $p < 0.001$, ns= non-significant.

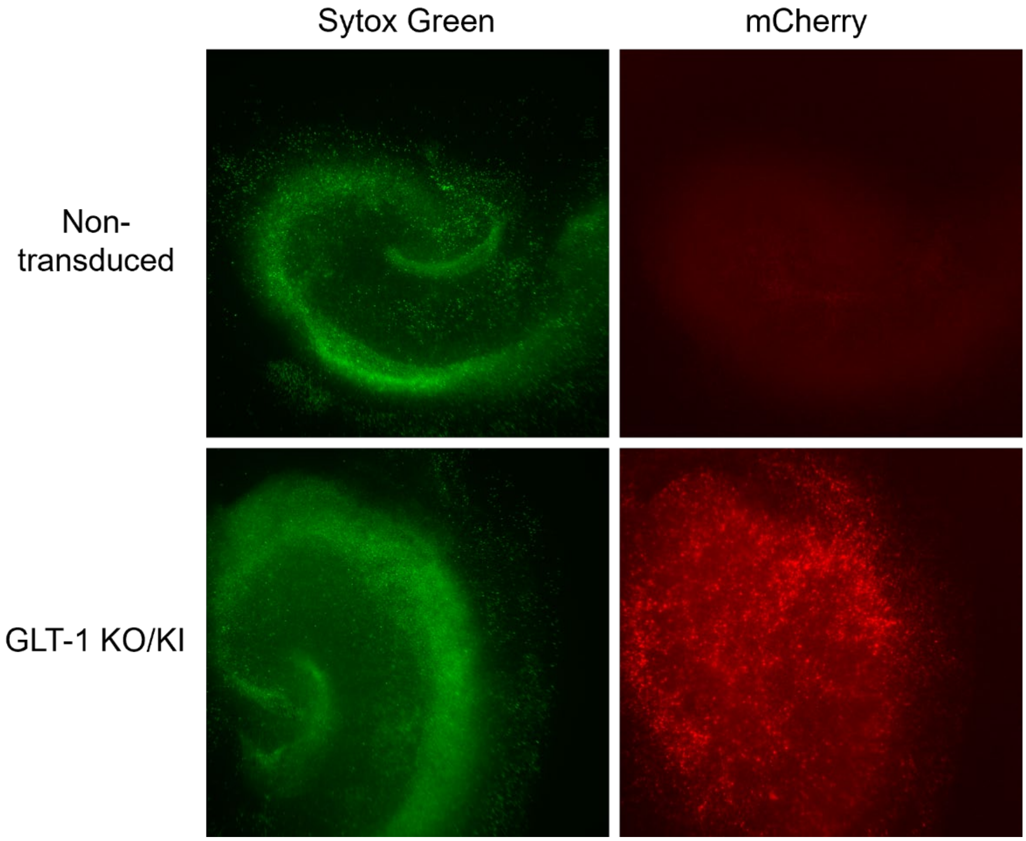


**Supplemental Figure 8. Validation of lentiviral transduction in hippocampal organotypic slice cultures (OSCs).** Representative immunofluorescence images of hippocampal OCSs transduced with GLT-1 KO/KI WT, compared to non-transduced slices, following OGD and Sytox Green staining. mCherry signal (red fluorescence) indicates successful viral transduction within the slice cultures.
